# Preserved respiratory chain capacity and physiology in mice with profoundly reduced levels of mitochondrial respirasomes

**DOI:** 10.1101/2023.06.19.545560

**Authors:** Dusanka Milenkovic, Jelena Misic, Johannes F Hevler, Thibaut Molinié, Injae Chung, Ilian Atanassov, Xinping Li, Roberta Filograna, Andrea Mesaros, Arnaud Mourier, Albert J R Heck, Judy Hirst, Nils-Göran Larsson

## Abstract

The mammalian respiratory chain complexes I, III_2_ and IV (CI, CIII_2_ and CIV) are critical for cellular bioenergetics and form a stable assembly, the respirasome (CI-CIII_2_-CIV), that is biochemically and structurally well documented. The role of the respirasome in bioenergetics and regulation of metabolism is subject to intense debate and is difficult to study because the individual respiratory chain complexes coexist together with high levels of respirasomes. To critically investigate the *in vivo* role of the respirasome, we generated homozygous knock-in mice that have normal levels of respiratory chain complexes but profoundly decreased levels of respirasomes. Surprisingly, the mutant mice are healthy, with preserved respiratory chain capacity and normal exercise performance. Our findings show that high levels of respirasomes are dispensable for maintaining bioenergetics and physiology in the mouse, but raises questions about their alternate functions, such as relating to regulation of protein stability and prevention of age-associated protein aggregation.

## INTRODUCTION

Respiratory chain supercomplexes were originally observed by BN-PAGE analyses following solubilization of mitochondrial membranes with mild ionic detergents, but their existence *in vivo* remained controversial for a long time ^1^. More recently, studies using a multitude of independent approaches, including electron cryo-microscopy (cryo-EM), electron cryo-tomography (cryo-ET), cross-linking mass spectrometry (XL-MS), and Förster resonance energy transfer (FRET) analyses, have convincingly proven their existence in various organisms ^2–8^. In particular, the respirasome (CI-CIII_2_-CIV) has attracted a lot of interest because it is the simplest possible supercomplex that contains a complete respiratory chain, wherein electrons enter from NADH at CI, are translocated by coenzyme Q, (Q) to CIII_2_, thereafter transferred to cytochrome *c* (Cyt *c*) and finally delivered to CIV to reduce O_2_ to H_2_O. It has also been reported that supercomplexes are dynamic and can be deconstructed, rebuilt and reconfigured to alter flux pathways depending on the metabolic conditions and available substrates ^9,10^. Isolated respirasomes are capable of functioning as independent bioenergetic units as they sequester their own dedicated Q and Cyt *c* molecules to transfer electrons between the complexes ^11^. Similarly, respirasomes in the membrane have also been proposed to sequester their own dedicated Q and Cyt *c* pools, but analyses of their structures, as well as of kinetic and spectroscopic data, do not support strict substrate channeling mechanisms. Instead, although kinetic advantages may result from binding site proximities, Q is able to exchange freely between complexes both in and out of the respirasome, and the positively-charged Cyt *c* moves in 2D sliding modes across the negatively-charged surfaces of CIII_2_ and CIV ^6,7,12–18^. The roles of respirasomes in bioenergetics therefore remain in question.

Aberrant respirasome levels have been reported in various types of human pathologies, including heart failure, ageing and age-related neurodegenerative disorders ^19–23^. There are also several reports arguing that increased formation of respiratory chain supercomplexes promote the growth of different types of cancer cells by enhancing hypoxia tolerance ^24–26^. Respiratory chain supercomplexes have also been reported to provide protection against influenza virus infection ^20^. An increase in the steady-state levels of respiratory supercomplexes have been found to improve mitochondrial bioenergetics and increase exercise capacity ^9,27,28^. In several studies of the role of respirasomes in physiology and disease, a rather modest change in respirasome levels have been reported to have marked effects on physiology or disease progression, e.g., ∼40% decrease in cerebral cortex in ageing rats ^23^, ∼30% decrease in heart of dogs with heart failure ^21^, ∼20-30% decrease in lymphoblasts in TAFAZZIN-deficient Barth syndrome patients ^22^ and ∼30-40% decrease in pancreatic cancer cells ^26^. A major problem with this type of studies is that levels of supercomplexes may correlate with steady-state levels of individual respiratory chain complexes. Altered levels of respirasomes or other types of supercomplexes in physiology or pathology may therefore just be a reflection of the degree of mitochondrial biogenesis.

One approach to probe the bioenergetic and physiological roles of the mammalian respirasome is to compare the properties of two otherwise-identical *in vivo* systems, one in which the individual complexes are stably bound to each other in respirasomes, and the other in which they are not. Important criteria for creating such a system would be that the individual complexes are fully assembled and functional, but that they do not form stable inter-enzyme interactions to form respirasomes (which we define as stable entities, not only the result of enzyme proximities). Here we have enabled such a comparison by generating a mouse model with a profound reduction in the levels of respirasomes, for comparison with the wild-type mouse that contains a normal and substantial complement of respirasomes. Analysis of mammalian respirasome structures identified three putative protein-protein interaction sites that could stabilize interactions by CI and CIII_2_ in the respirasome. By generating mouse knock-in strains, we discovered that one of the putative interactions between CI and CIII_2_ was critical. Loss of three charged amino acids (EED) in the UQCRC1 subunit of CIII_2_ drastically decreased the interaction between CI and CIII_2,_ leading to a profound decrease of stable respirasomes on BN-PAGE gels in all investigated tissues. Surprisingly, comprehensive evaluation of the effects of the loss of stable respirasomes revealed a maintained respiratory chain function and normal motor performance during exercise. This is surprising in the light of how essential respirasomes are considered to be for the function of the bioenergetic membrane, and also because XL-MS data support that the organization of respiratory chain complexes in the membrane is perturbed. The mouse model with a drastic decrease of respirasomes that we have developed here will be a valuable tool for future studies of the role of respirasomes in physiology, disease, and ageing.

## RESULTS

### Disruption of critical protein-protein contacts in the respirasome

To work out how to disrupt the interactions between the respiratory chain complexes that form the respirasome, we analyzed available respirasome structures to predict critical protein-protein interaction sites between CI and CIII_2_ and created three knock-in mouse models (Figures 1A-C and S1A-C). The aim was to remove the stabilizing interactions that hold the complexes together in the specific, structurally homogeneous, stable assembly that is defined as the respirasome. Given the substantial literature highlighting the importance of respirasomes for proper function of the oxidative phosphorylation (OXPHOS) system ^29,30^, we used conditional knock-in strategies to circumvent the possibility of embryonic lethal phenotypes (Figures S1A-C). In the first strain, a conserved tyrosine was mutated to an alanine in the NDUFA11 subunit (*Ndufa11^Y110A^*) of CI that contacts the UQCRQ subunit of CIII_2_ (Figure 1A). This conditional knock-in allele was established by introducing loxP sites around the last exon of *Ndufa11*, followed by a mutant version of the exon encoding the Y110A substitution (Figure S1A). In the second strain, the C-terminal 20 amino acids (K118-L137) of CI subunit NDUFB7 (*Ndufb7^DEL: K118-L137^*) were deleted (Figure 1B). The C-terminal helix of NDUFB7 extends away from CI in a helix but then is not observed clearly in structural data; it may form interactions with CIII_2_ and/or CIV (Figure 1B). This conditional knock-in allele was generated by flanking exon 2 and 3 of *Ndufb7* with loxP sites, followed by a duplication of exons 2 and 3 in which exon 3 has the introduced deletion (Figure S1B). In the third strain, a short loop of three charged amino acids was deleted from the UQCRC1 subunit of CIII_2_ (*Uqcrc1^DEL:E258-D260^*) as they are observed to interact with the NDUFB4 and NDUFB9 subunits of CI (Figure 1C). Interestingly, this short loop is not present in the UQCRC1 subunit of *Saccharomyces cerevisiae*, which does not contain CI. The conditional knock-in allele for *Uqcrc1* was generated by introducing loxP sites flanking exons 7–13, followed by a duplicated genomic region of exon 7–13 in which nine nucleotides are deleted from exon 7 (Figure S1C). We proceeded to breed the mice to generate strains that were homozygous for the loxP-flanked knock-in alleles and heterozygous for a transgene expressing cre-recombinase under the control of the muscle creatine kinase promoter (Ckmm-cre), which is expected to activate the variant alleles in heart and skeletal muscle. The expected genomic rearrangements were confirmed by PCR genotyping followed by sequencing.

**Figure 1.**
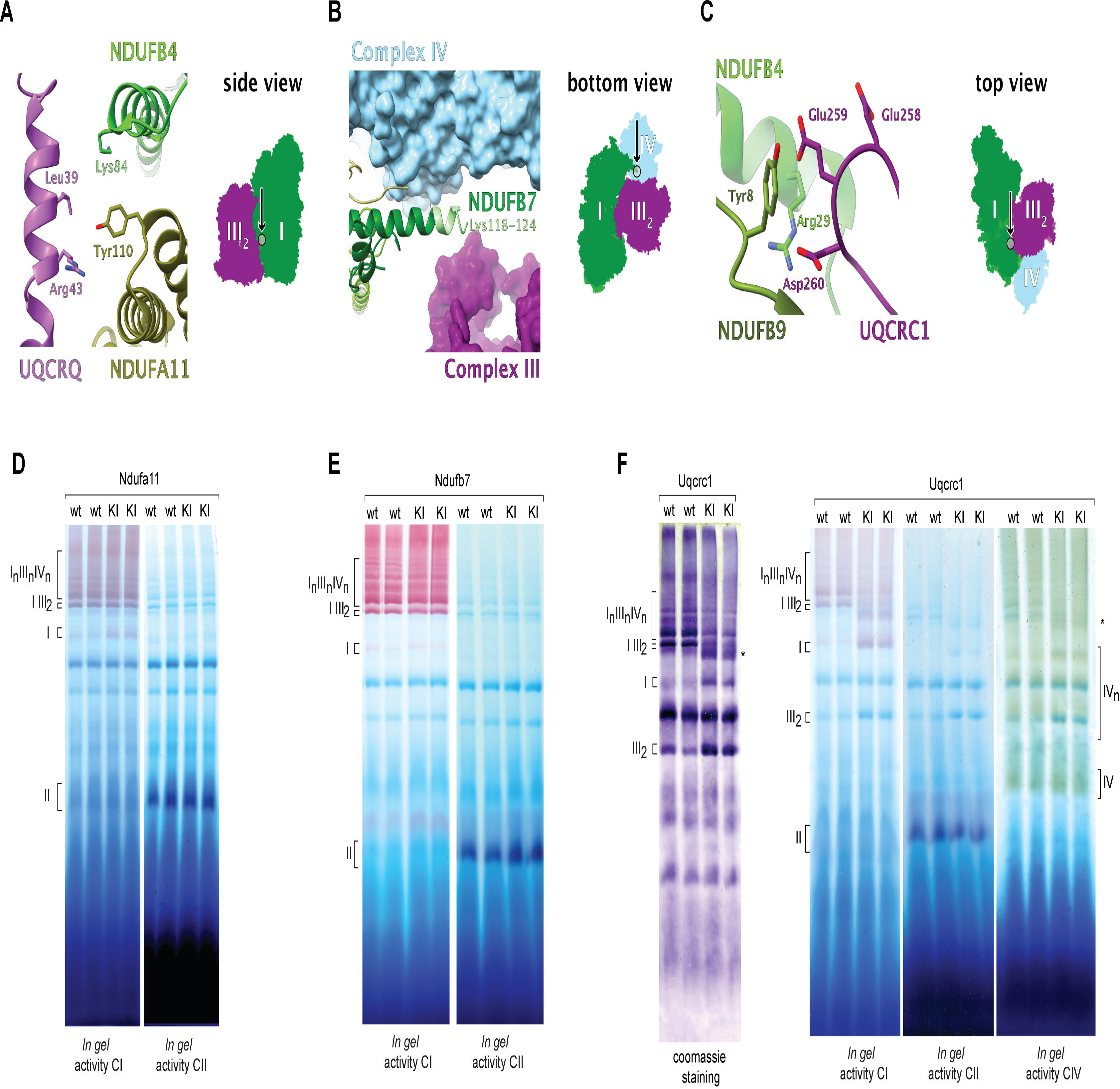
Mutations at three structurally-defined respirasome contact sites. (A-C). The structurally-defined interactions between different respiratory chain complexes in respirasomes that were disrupted (PDB: 5GUP ^33^ and 6QBX ^61^; the models were mutated in ChimeraX ^62^ to reflect mouse-specific residues) (D-F) The effects of the *Ndufa11^Y110A^, Ndufb7^DEl:K118-L137^ and Uqcrc1^DEL:E258-D260^* knock-in mutations on supercomplex organization in heart mitochondria of tissue specific mutants analyzed by BN-PAGE followed by Coomassie staining or CI, CII and CIV *in gel* activity assays. The CII in gel activity serves as a loading control. * novel supercomplex species formed upon introduction of the Uqcrc1*^DEL:E258-D260^* knock-in mutation.

In BN-PAGE analyses of heart mitochondria of the tissue-specific knock-in mice, the *Ndufa11^Y110A^* variant exhibited mildly decreased levels of respirasomes and slightly increased levels of free CI (Figure 1D). Harsher solubilization conditions with a higher digitonin to protein ratio did not cause further destabilization of respirasomes (Figure S2A). No obvious effects on supercomplex formation were observed in the tissue-specific *Ndufb7^DEL:K118-L137^* knock-in mice (Figure 1E) even when using stringent detergent solubilization conditions (Figure S2B). Western blots confirmed expression of the truncated NDUFB7 protein (Figure S2C). In contrast to the other two mutants, the tissue-specific *Uqcrc1^DEL:E258-D260^* knock-in mice (Figure 1C) exhibited drastically decreased levels of respirasomes, and clearly increased amounts of free CI and CIII_2_ (Figure 1F). Surprisingly, the mice from this strain also appeared healthy with no obvious phenotype.

### Mice expressing mutant UQCRC1 in all tissues have drastically decreased levels of respirasomes

As the tissue-specific *Uqcrc1^DEL:E258-D260^* mice, which have very low levels of stable respirasomes in heart, appeared healthy, we proceeded to activate the expression of the knock-in allele in all tissues by breeding conditional knock-in *Uqcrc1^DEL:E258-D260^* mice to mice ubiquitously expressing cre-recombinase from the *β-*actin promoter. Successful deletion of nine nucleotides from exon 7 of the *Uqcrc1* gene was confirmed by PCR (Figure S2D) and the *β-*actin-cre transgene was removed by breeding. An intercross of heterozygous mice generated offspring at the expected Mendelian ratios (for 336 offspring from 47 litters: 82 *Uqcrc1^+/+^* [24.4%]; 166 *Uqcrc1^+/DEL:E258-D260^* [49.4%]; 88 *Uqcrc1^DEL:E258-D260/DEL:E258-D260^* [26.2%]). The whole-body homozygous knock-in mice (*Uqcrc1^DEL:E258-D260/DEL:E258-D260^*; hereafter referred to as *Uqcrc1* knock-in mice) were viable and did not show any obvious phenotype, and were used for all subsequent experiments in this study. The *Uqcrc1* mutation did not affect the steady-state levels of UQCRC1 or other OXPHOS proteins in isolated mitochondria of heart, kidney, liver, skeletal muscle, and brain (Figure S2E). The *Uqcrc1* knock-in mice were fertile and produced normal litter sizes (mean 6.7 pups from ten litters of five independent matings).

Strikingly, BN-PAGE analyses of mitochondria isolated from different tissues of *Uqcrc1* knock-in mice revealed an absence of supercomplex I/III_2_, a substantial decrease in respirasome content and release of CI and CIII_2_ from supercomplexes to free pools (Figure 2A). Importantly, BN-PAGE analyses performed after solubilization of mitochondria by n-dodecyl-β-D-maltoside (DDM) showed that homozygosity for the *Uqcrc1* knock-in allele does not affect the assembly or stability of the individual respiratory chain complexes (Figure 2B) and tandem mass tag-based quantitative proteomics showed that the levels of a range of individual respiratory chain subunits were normal (Figure S2F). The decreased respirasome content is thus not caused by destabilization of the individual respiratory chain complexes.

**Figure 2.**
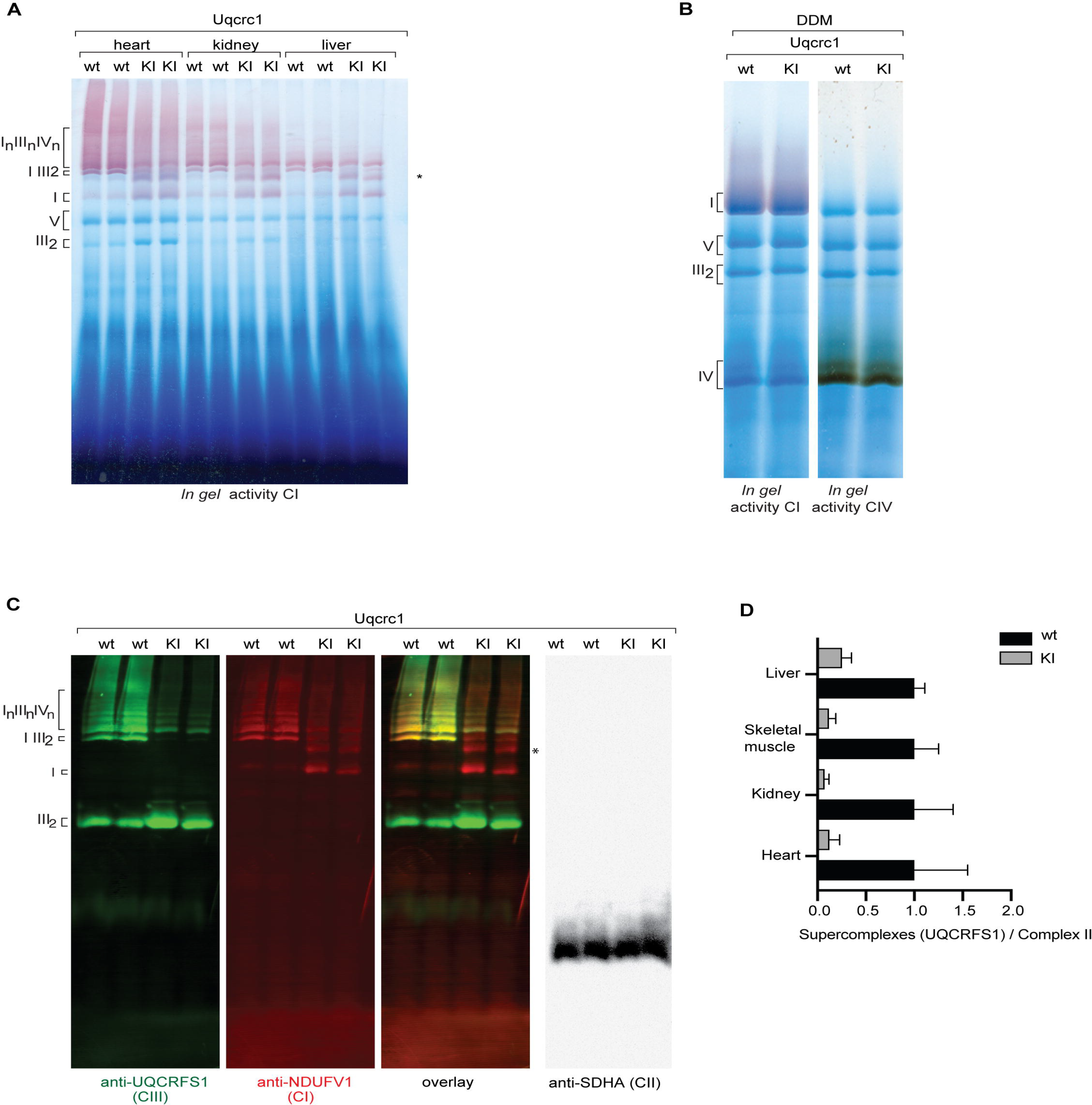
The *Uqcrc1^DEL:E258-D260^* mutation leads to a drastic reduction of stable respirasomes in various tissues of mice. (A) Complex I *in gel* activity of mitochondria isolated from various wild-type and *Uqcrc1^DEL:E258-D260^*-expressing tissues. (B) BN-PAGE analysis of wild-type and *Uqcrc1^DEL:E258-D260^* mitochondria after DDM solubilization. The analysis is followed by CI and CIV *in gel* activity assays. (C) Fluorescent western blot analyses of respiratory chain organization in wild-type and *Uqcrc1^DEL:E258-D260^* -expressing heart mitochondria. Indicated antibodies were used for detection of respiratory chain complexes. (D) Quantification of UQCRFS1/CIII signals in the molecular weight range corresponding to respirasome from wild-type and *Uqcrc1* knock-in mice. BN-PAGE signals from four different tissues were quantified using SDHA/CII as a reference. n=6 per group, error bars indicate mean ± SEM. Statistical test: unpaired Welch’s t test.

BN-PAGE analyses with Coomassie staining or CI *in gel* activity assays (Figures 1F and 2A) showed a marked reduction of supercomplexes in *Uqcrc1* knock-in mice and we therefore proceeded with fluorescent western blot analyses (Figure 2C). With this method we found a drastic decrease of co-migration between CI and CIII_2_ in the high-molecular weight fractions containing respirasomes in *Uqcrc1* knock-in mice (Figure 2C). The reduction in respirasomes was accompanied by increased levels of free CI and CIII_2_, and the appearance of a novel CI-containing supercomplex (Figure 2C). We proceeded to quantify supercomplexes by using BN-PAGE and western blots with an antibody against the catalytic core subunit of CIII_2_ (UQCRFS1). The levels of CIII_2_-containing supercomplexes were reduced by 75-90% in liver, skeletal muscle, kidney, and heart of *Uqcrc1* knock-in mice (Figure S2D), consistent with a profound depletion of respirasomes.

To further quantitatively assess the levels of respiratory chain subunits in supercomplexes, we performed complexome profiling proteomics of heart mitochondria resolved by BN-PAGE (Figures 3A-B and S3). We quantified the differences in OXPHOS complex abundance in BN-PAGE slices by using the MaxQuant’s MaxLFQ algorithm ^31^ and found a 80-90% depletion of CIII_2_ subunits in the supercomplex fractions, along with a substantial shift in the distribution of CI and CIII_2_ towards free enzymes (Figure 3B).

**Figure 3.**
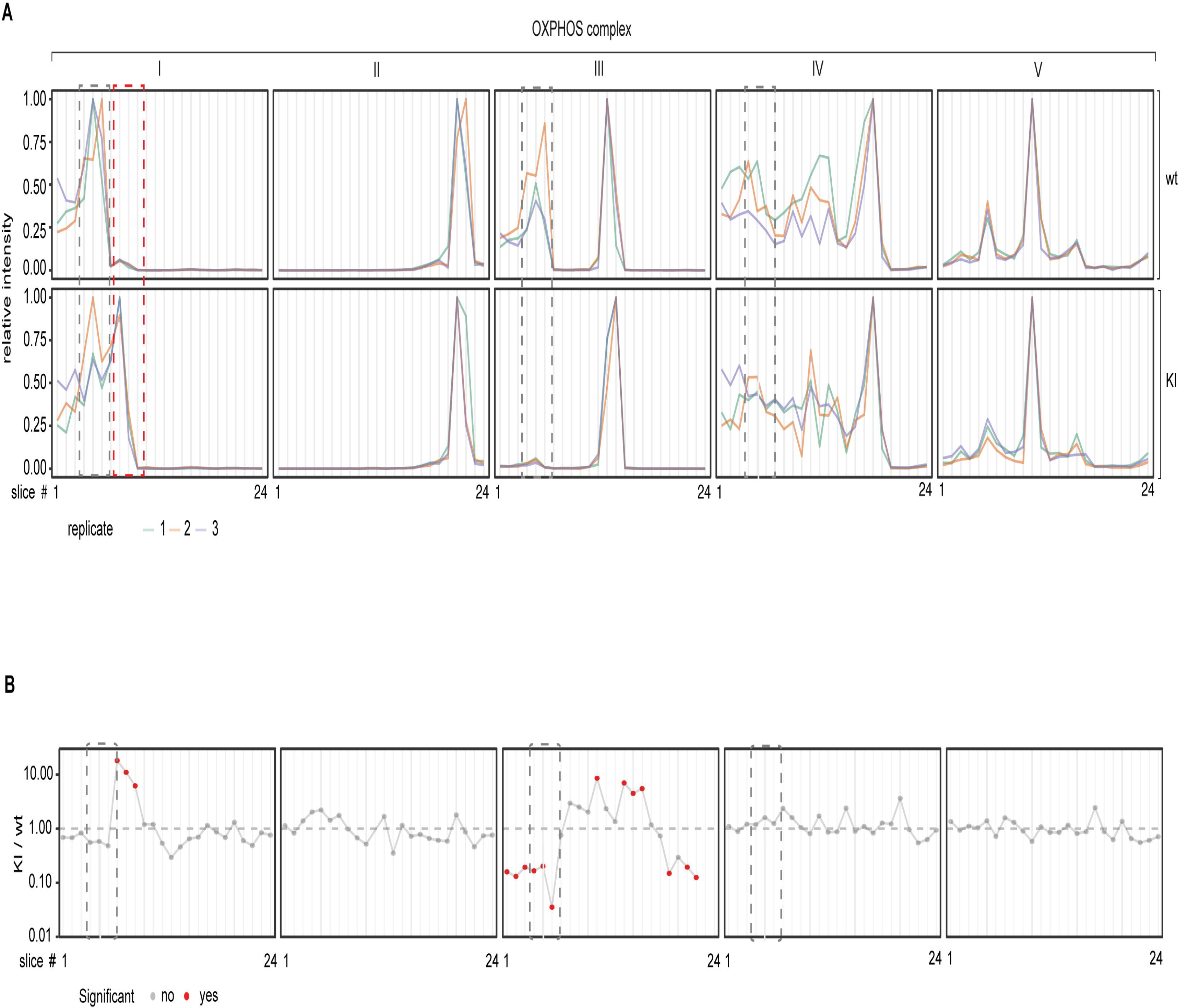
Complexome profiling of OXPHOS proteins from heart *Uqcrc1^DEL:E258-D260^* mitochondria and controls. (A) Distribution of OXPHOS complexes CI-CV along the BN-PAGE gel in wild-type and *Uqcrc1^DEL:E258-D260^* knock-in mice. Each profile represents the sum of intensities of all subunits of a complex, relative to the maximum intensity of that complex in the respective BN-PAGE lane. In *Uqcrc1^DEL:E258-D260^* knock-in mice, a novel CI-containing supercomplex species that lacks CIII_2_ is indicated by the red hatched box, whereas the position of supercomplex I/III_2_ and the respirasome is indicated by the grey hatched box. (B) Differences in abundance between KI and WT per BN-PAGE slice. The intensities of all subunits were added up to derive the intensities of the whole complexes that were used for differential abundance analysis; the position of supercomplex I/III_2_ and respirasome is indicated by the grey hatched box. Please note that the y axis is logarithmic.

Furthermore, a novel, high-molecular weight supercomplex, migrating above free CI (Figures 1F, 2A and 2C; marked with *), was present in the *Uqcrc1* knock-in mitochondria. *In gel* activity assays (Figures 1F and 2A) and fluorescent western blot analysis (Figure 2C) suggested that this novel supercomplex is mainly composed of CI subunits. To further assess the composition of the novel complex that forms in response to the UQCRC1 microdeletion, we performed quantitative complexome analyses of heart, skeletal muscle and kidney from *Uqcrc1* knock-in mice. From each tissue, we analyzed the area containing the novel aberrant complex (slice 3 from the BN gel) as well as neighbouring regions containing higher molecular complexes (slice 1 and 2) and lower molecular complexes (slice 4 and 5). This comprehensive analysis shows that the novel aberrant complex (fraction 3) contains significantly enriched CI subunits in all three tissues and no other proteins (Figure S4). This observation suggests that ablation of the specific stabilizing interaction between CI and CIII_2_ in the respirasome has allowed the formation of new interactions and thus the formation of new complex assemblies that nonetheless cannot functionally replace the respirasomes.

### Mice with very low levels of respirasomes have normal bioenergetic capacity

To analyze the bioenergetic properties of the mutant mouse mitochondria, we performed kinetic activity assays on mitochondrial membranes isolated from wild-type and *Uqcrc1* knock-in hearts. BN-PAGE analyses of the OXPHOS complexes in mitochondrial membranes (Figure S5A) confirmed the previous findings from whole mitochondria (Figures 1F, 2A and 2C) and showed a reduction of supercomplexes containing CIII_2_. Despite the drastic decrease in respirasomes, the rates of NADH oxidation catalyzed by CI, CIII_2_ and CIV, and succinate oxidation catalyzed by CII, CIII_2_ and CIV (Figures 4A and 4B) as well as NADH-induced reactive oxygen species (ROS) production by CI (Figure 4C) were not altered in *Uqcrc1* knock-in mice in comparison to wild-type controls. Moreover, we permeabilized mitochondria to specifically measure respiratory chain activity in mitochondria fed directly with CI (NADH) or CII (succinate) substrates or both substrates simultaneously and found no difference in respiration capacities between *Uqcrc1* knock-in and wild-type heart (Figure 4D). Furthermore, the oxygen consumption rates (Figure 4E) and the hydrogen peroxide production (Figure S5B) with complex I or complex II substrates under phosphorylating, non-phosphorylating, and uncoupled conditions were all similar in *Uqcrc1* knock-in and wild-type mouse heart mitochondria. We also assessed OXPHOS yield (JATP/JO_2_) and found no difference between wild-type and knock-in mitochondria (Figure S5C), suggesting that decreased respirasome levels do not influence the coupling between respiration and ATP synthesis and that proton pumping by the respiratory chain is unchanged under the studied conditions.

**Figure 4.**
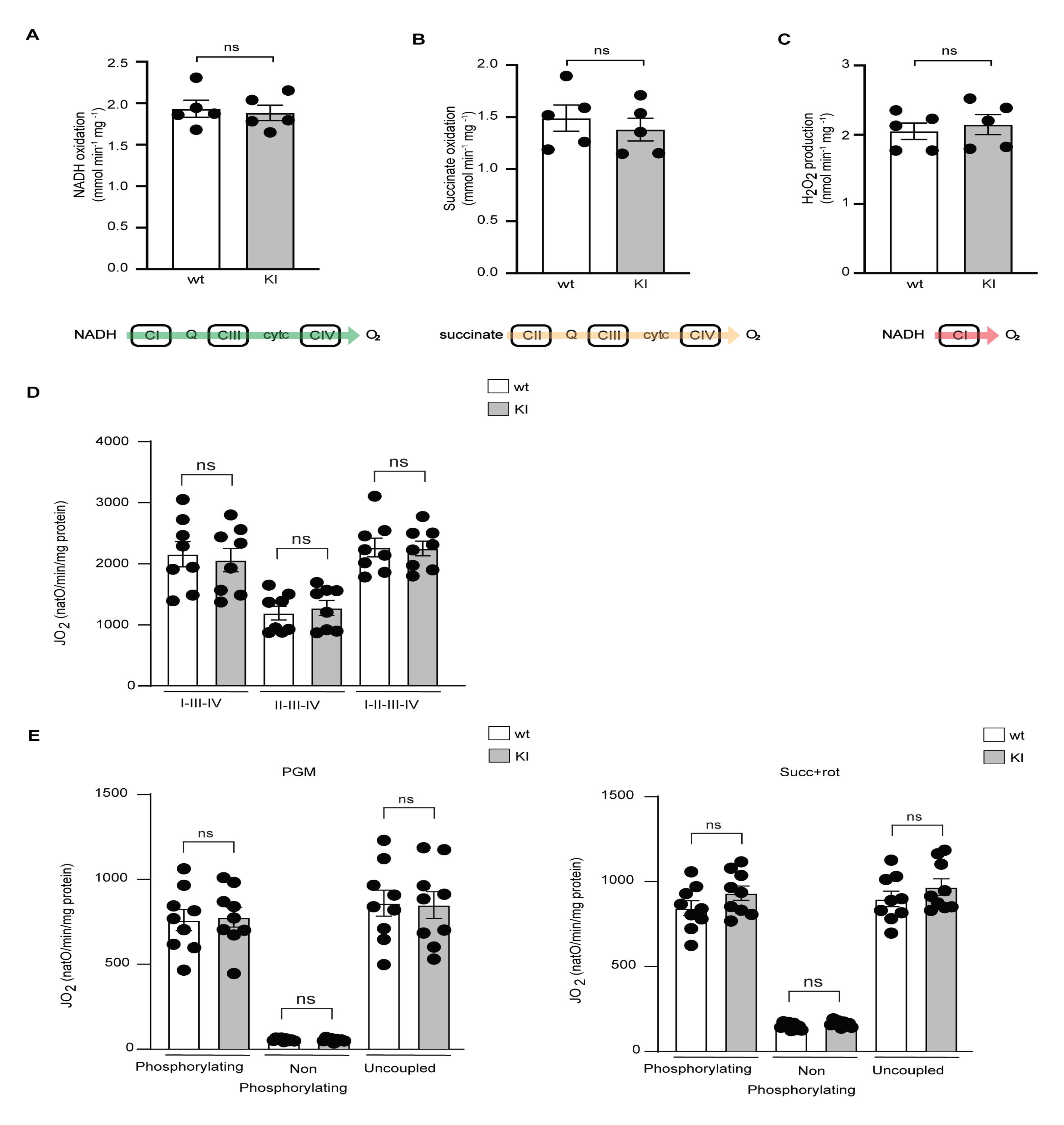
Normal bioenergetic capacity and ROS production in mice with very low levels of stable respirasomes. (A and B) NADH:O_2_ and succinate:O_2_ oxidoreductase activities of mitochondrial membranes of wild-type (white bars) and *Uqcrc1* knock-in (grey bars) mice. Reported values are the mean averages from n = 5 membrane preparations (each from different hearts) and are represented as ± SEM values. Statistical test: unpaired students t test (two-tailed). (C) NADH-stimulated H_2_O_2_ production by CI in wild-type (white bars) and *Uqcrc1* knock-in (grey bars) mitochondrial membranes. n = 5; error bars indicate mean ± SEM. Statistical test: unpaired students t-test (two-tailed). (D) The maximal oxygen consumption rate assessed in permeabilized heart mitochondria from wild-type (white bars) and *Uqcrc1* knock-in (grey bars) mice in the presence of non-limiting concentrations of substrates. n = 8; error bars indicate mean ± SEM. Statistical test: one-way ANOVA. (E) Oxygen consumption of heart mitochondria from wild-type (white bars) and *Uqcrc1* knock-in (grey bars) mouse strains at 22 weeks of age. n = 9; error bars indicate mean ± SEM. Statistical test: one-way ANOVA. Abbreviations: PGM, pyruvate, glutamate and malate; Succ, succinate; rot, rotenone.

In line with data obtained from heart, the respiration capacity in liver mitochondria of *Uqcrc1* knock-in mice was also unaffected (Figure S6A). The oxygen consumption rate with complex I (Figure S6B) or complex II (Figure S6C) substrates under phosphorylating, non-phosphorylating, and uncoupled conditions were also similar in *Uqcrc1* knock-in and wild-type mouse liver mitochondria. These results are at variance with other reports arguing that supercomplexes are necessary to promote optimal and simultaneous oxidation of different substrates^9^. Moreover, the enzyme activities of individual (CI, CII and CIV) or combined complexes (CI-CIII and CII-CIII) were not affected by the *Uqcrc1* mutation in heart and liver mitochondria (Figures S6D and S6E).

### Mice with very low levels of respirasomes have normal motor activity and exercise performance

We proceeded with phenotypic characterization of *Uqcrc1* knock-in and wild-type mice with a special focus on locomotor activity and exercise performance (Figures 5 and S7). Wild-type and mutant animals of both genders at age 25 weeks performed equally well in a RotaRod test (Figures 5A and 5B) and evaluation of the exploratory locomotor activity showed no differences between the groups when total distance (Figure 5C), total moving time (Figure 5D), overall speed (Figure 5E), and time spent in the periphery (Figure 5F) were assessed. To evaluate exercise performance, cohorts of *Uqcrc1* knock-in and wild-type mice of both sexes (16 males and 8 females per genotype in three independent experiments) were trained by allowing access to a running wheel situated in the cage for fifteen days. In addition, three treadmill running sessions were performed at the beginning (day 1), day 8 and at the end (day 14) of the experiment (Figure S7A). Body weight and fat content of the mice were assessed at day 1 and 14 of the experiment and did not differ significantly between the genotypes (Figures S7B and S7C). The male and female animals of both genotypes showed a tendency for decreased fat content upon exercise (Figure S7C). Importantly, *Uqcrc1* knock-in and wild-type mice were running comparable distances in both forced exercise experiments on a treadmill (Figure S7D) and voluntary exercise on a running wheel situated in the cage (Figure S7E) thus showing that the profound decrease of respirasomes does not affect exercise capacity.

**Figure 5.**
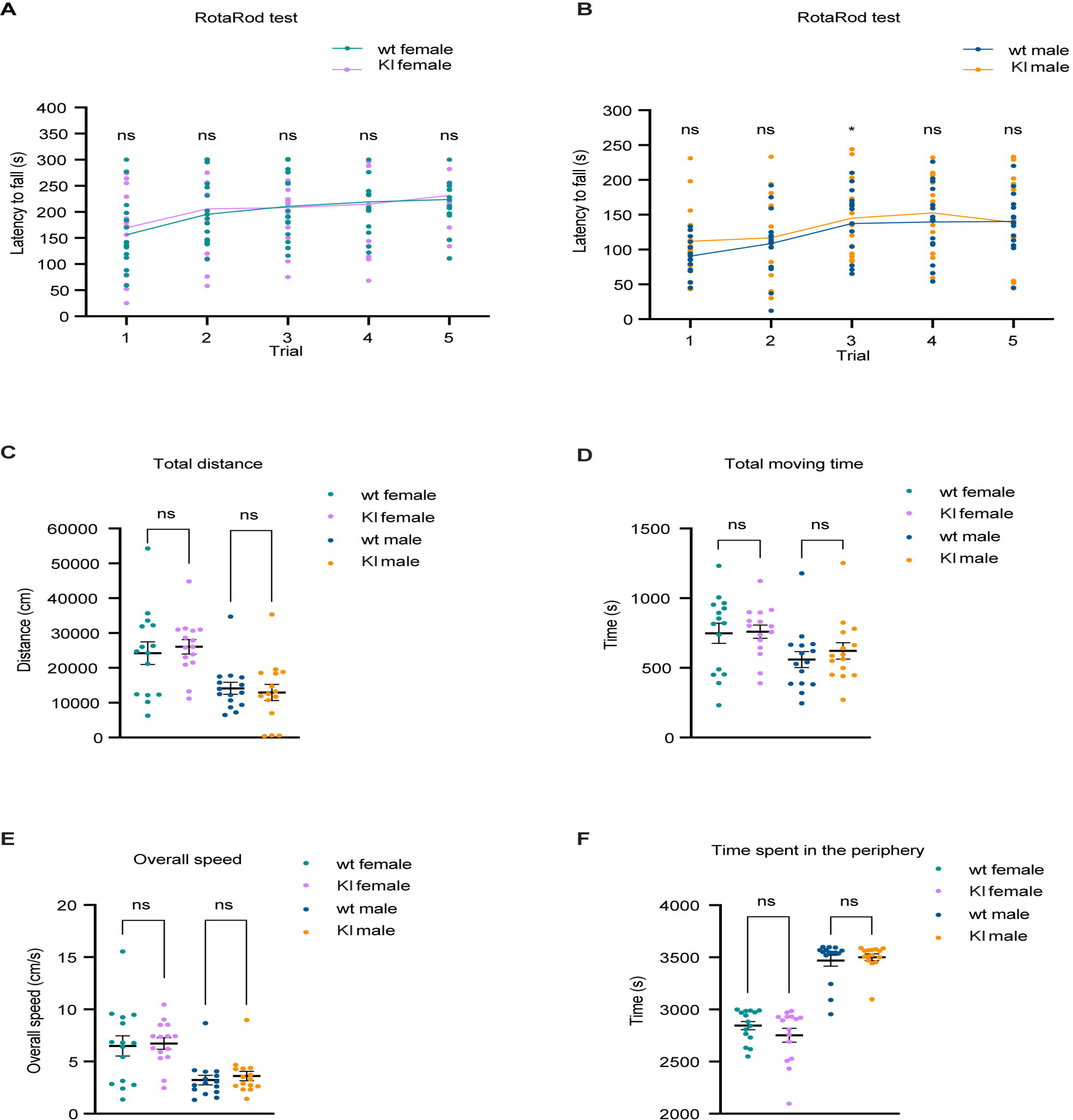
Normal motor activity in mice with very low levels of stable respirasomes. (A and B) RotaRod test was performed with wild-type and *Uqcrc1* knock-in mice of both sexes for five successive trials. (C-F) Open field experimental test was carried out with wild-type and *Uqcrc1* knock-in mice of both genders and total distance (C), total moving time (D) overall speed (E) and time spent in the periphery (F), were recorded. A total of 60 animals of both genotypes and genders at 25 weeks of age were used for the experiment. Statistical analyses were performed using Welch’s t-test. Error bars indicate mean ± SEM.

To address putative metabolic differences, we collected data for 48 hours from control and *Uqcrc1* knock-in mice of both sexes in metabolic cages. We did not observe any consistent changes in total activity, food and water intake, VO2, VCO2 and the respiratory exchange ratio (RER) between the two genotypes (Figures S8A - S8L). Therefore, the decrease in stable respirasomes does not obviously affect the metabolic parameters of mice analyzed in metabolic cages.

### Cross-linking of intact mitochondria shows modulated respiratory chain organization in *Uqcrc1* knock-in mice

The finding of substantially decreased levels of respirasomes by BN-PAGE analyses and complexome proteomics profiling (Figures 1F, 2A, 2C, 3A and 3B) accompanied by maintained respiratory chain function (Figures 4 and S5B, S5C, S6A, S6B and S6C) and normal motor performance (Figures 5 and S7) in *Uqcrc1* knock-in mice is unexpected. These findings contradict literature reports arguing that modest changes in respirasome levels or supercomplex organization have a marked impact on bioenergetics and physiology ^9,26–28^. Our results showing a marked decrease of stable respirasomes are based on treatment of mitochondria with mild detergent, which is the standard procedure for analysis of respirasomes with BN-PAGE ^1,32^ and for isolation of respirasomes for single particle cryo-EM structure determination ^6,7,33^. To exclude that the observation of decreased respirasomes in *Uqcrc1* knock-in mice is fostered by solubilization and BN-PAGE separation, we next aimed at stabilizing mitochondrial complexes by cross-linking mitochondria prior to membrane solubilization and BN-PAGE separation using DMTMM as cross-linking reagent ^34^ (Figure 6A). Importantly, BN-PAGE analysis of DMTMM cross-linked mitochondria corroborate our previous findings (Figures 1F, 2A, 2C, 3A and 3B), suggesting that neither solubilization nor BN-PAGE analysis artificially promote respirasome disruption in *Uqcrc1* knock-in mice.

**Figure 6.**
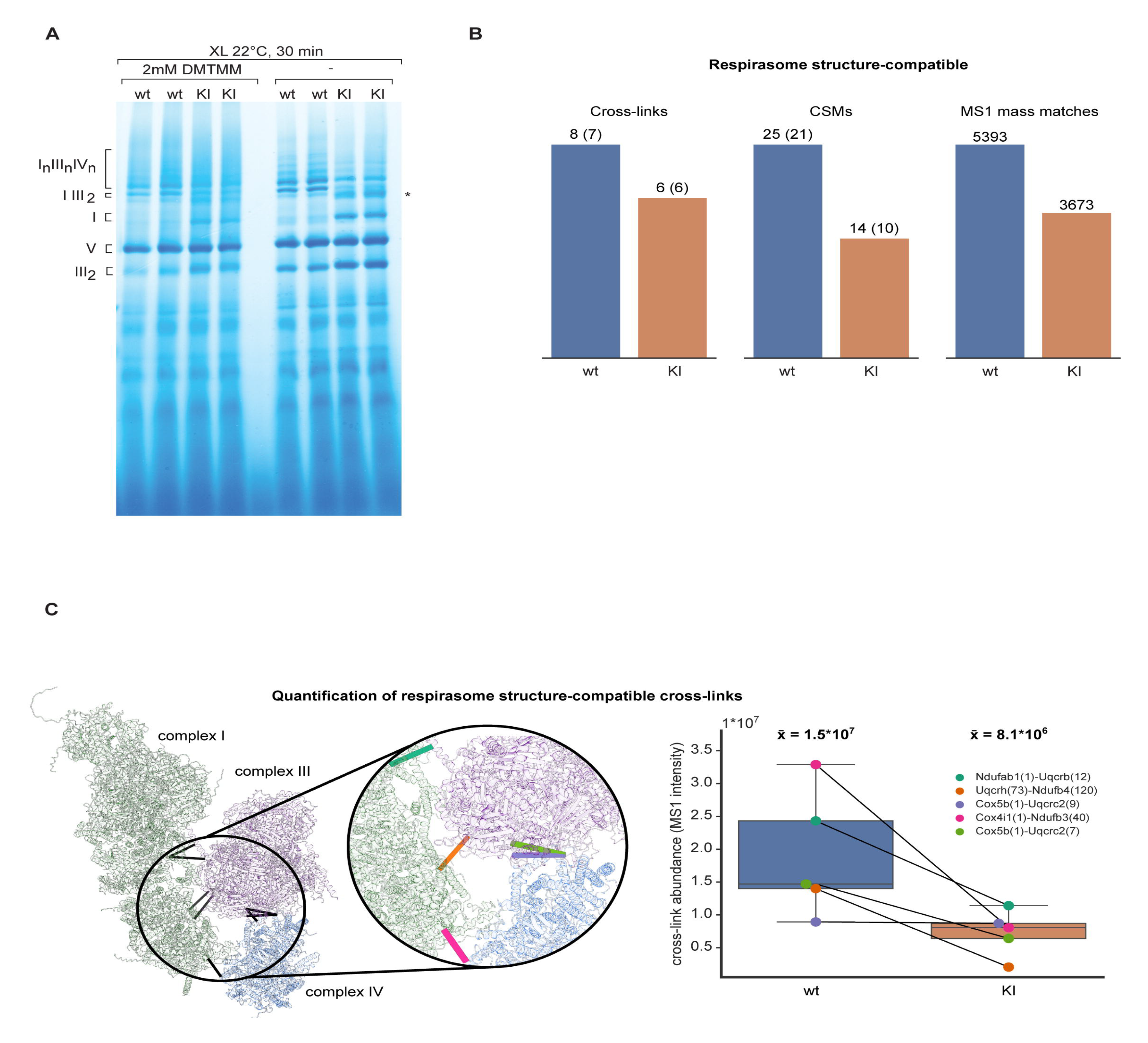
Cross-link frequencies and intensities involving respirasomes in wild-type and *Uqcrc1* knock-in mitochondria. (A) BN-PAGE analysis of wild-type and *Uqcrc1* knock-in mitochondria upon DMTMM crosslinking. (B) Count of respirasome structure-compatible cross-links (distance < 30 Å), underlying cross-link spectral matches (CSMs) and MS1 mass matches in wild-type (blue) and Uqcrc1 knock-in (orange) replicates. MS1 mass matching of identified cross-linked peptide precursors was performed with a mass tolerance of +/- 30mDa. The number of cross-links and CSMs with an identified precursor abundance (determining whether an observed cross-link can be quantified) is indicated in brackets. (C) Structural model of the mouse respirasome. Previously resolved structures for Complex I (green), Complex III (purple) and Complex IV (blue) (PDB: 6ZR2,7O3H, 7O3E) were structurally aligned based on the Sus scrofa respirasome structure (PDB: 5GUP). Where applicable, cross-linked, previously unresolved N-termini were modelled using Alphafold. Cross-links with a distance < 30 Å observed for wild-type and Uqcrc1 knock-in were mapped onto the structure and classified as respirasome structure-compatible (black, solid lines). Quantified cross-links that are observed for both wild-type (wt) and knock-in (KI) mitochondria are shown. Each colour represents a specific cross-link (left and right panel). The right panel with box plots shows the abundance of the individual cross-links in wild-type (blue) and Uqcrc1 knock-in (orange) mitochondria. Each black line in the right panel indicates the individual abundance change for a specific cross-link observed in both wild-type and knock-in mitochondria.

Aspiring to further probe the *in vivo* spatial arrangement of respiratory chain proteins in wild-type and *Uqcrc1* knock-in mice, we performed cross-linking mass-spectrometry (XL-MS) on intact mitochondria using an optimal concentration of lysine reactive reagent disuccinimidyl sulfoxide (DSSO, Figure S9A) ^3,35–38^. On proteome-wide scale, no significant differences between conditions (Figures S9B) and good reproducibility were observed (Figure S9C and Table S1). Next, we specifically analyzed cross-links for the respiratory chain proteins I, III and IV. In line with previous studies ^34,36–38^, cross-links of respirasome complexes were exclusively found for solvent accessible domains, frequently involving their highly exposed and mobile N-termini. Given that commonly available cross-linkers such as DSSO are not capable of probing protein-protein interactions within the membrane, we first set out to generate a more complete model of the solvent accessible parts of the respirasome. Mouse CI (PDB 6ZR2), CIII (PDB 7O3H) and CIV (PDB 7O3E) structures were aligned to the structure of the respirasome from *Sus scrofa* (5GUP). The generated extended model is in good agreement with our cross-linking data (Figure S9D), for which mapped intra-protein cross-links are predominately below the DSSO distance threshold of 30 Å (x̃_*complex I*_ = 16.6 Å, x̃_*complex III*_ = 18.0 Å, x̃_*complex IV*_ = 26.8 Å) (Figure S9D).

To identify cross-links reflective of an assembled respirasome, we next set out to classify the observed cross-links. Inter complex cross-links with a distance <= 30 Å were classified as respirasome structure-compatible (Figure 6B). For wild-type and *Uqcrc1* knock-in mice, we identified eight and six cross-links, respectively, supported by identification of 25 and 14 cross-link spectral matches (CSMs) (Table S1). As the cross-link identification efficiency suffers from the low abundant nature of cross-linked peptides^39^, we performed MS1 mass matching, aiming at detecting cross-link peptides that are present but have not been identified by the MS acquisition algorithm^40^. Overall, no significant differences for recorded MS1 mass tags were observed (Figure S9E) between samples. However, masses corresponding to previously identified respirasome structure compatible cross-links are more frequently found in wild-type (5393) than in *UQCRC1* knock-in mice (3673) (Figure 6B). To further strengthen the observation of profoundly reduced levels of respirasomes in *UQCRC1* knock-in mice, we next set out to quantify identified cross-links by combining precursor intensity with spectral counts. On a proteome-wide scale 1728 out of 2074 (wild-type) and 1711 out of 2089 (knock-in) cross-links could be quantified, showing no significant abundance difference between genotypes (Figure S9B). Likewise, intra respirasome cross-links show only a slight difference in abundance, with cross-links observed in *UQCRC1* knock-in mice being slightly more abundant than intra links observed in wild-type (median abundance of 9.7*10^6^ vs 8.5*10^6^) (Figure S9F). On the contrary, quantifiable respirasome compatible cross-links showed a drastic abundance decrease (> 50 %) in *UQCRC1* knock-in mice compared to the wild-type (Figure 6C).

It is worth mentioning that besides respirasome compatible cross-links, we also identified respirasome-structure incompatible cross-links (distance > 30 Å). However, for these cross-links we found no significant differences regarding frequency and abundance between the genotypes (Table S1). Considering the architecture of the inner mitochondrial membrane and preferential localization of the respiratory complexes in the cristae ^41^, incompatible cross-links may hint at a proximity of individual complexes residing in opposing membranes or reflect random proximity effects due to the high protein density within the inner membrane.

## DISCUSSION

We demonstrate here that mice with profoundly reduced levels of respirasomes according to BN-PAGE analyses have maintained respiratory chain function and are healthy with normal motor activity and exercise capacity, thereby challenging the proposed requirement of stable, structurally defined respirasomes for maintaining optimal mitochondrial function ^9,10,28^. Consistent with our results, it was recently reported that destabilization of the interaction between CI and CIII_2_ does not have major consequences for plant physiology ^42^.

Supercomplexes containing CIII_2_ and CIV are observed in a wide range of bacteria and mitochondria of different eukaryotes. There is a remarkable difference in composition and architecture of the components amongst CIII_2_-CIV_1/2_ supercomplexes of different organisms and the only common structural feature seems to be the physical proximity of CIII_2_ and CIV ^43^. *In vitro* experimental evidence in budding yeast suggests that this proximity facilitates electron transfer by promoting 2D diffusion of Cyt *c* along the surface of the CIII_2_-CIV_1/2_ supercomplex ^17,43^. There has been a lot of controversy surrounding the role of the CIII_2_-CIV supercomplex in mammals. Whereas some studies have argued that the lowly abundant CIII_2_-CIV supercomplex, whose formation is mediated by COX7A2L, is linked to the formation of respirasomes and thereby plays an important role in bioenergetics and physiology ^9,44^, other studies have reported that the absence of the CIII_2_-CIV supercomplex does not affect respirasome formation or bioenergetics in human cell lines ^45,46^ or mice ^47,48^. A recent study reports that genetic variation affects the expression of human COX7A2L and thereby supercomplex formation and respiration in differentiated human myotubes ^49^. Because exercise induces mitochondrial biogenesis in skeletal muscle, it is difficult to distinguish whether training-induced increase of respirasome levels are due to specific effects on supercomplex formation or just reflect an increased abundance of respiratory chain complexes. A recent comprehensive study that assessed training-induced changes in human skeletal muscle by using multiple omics approaches reported that increased mitochondrial respiration induced by exercise is mediated by increased mitochondrial biogenesis without accompanying changes in the abundance and/or organization of respiratory supercomplexes in individual mitochondria ^50^.

The structure of supercomplex CIII_2_-CIV shows that the COX7A2L subunit is deeply integrated into both enzyme complexes and promotes a stable interaction between CIII_2_ and CIV ^51^. In contrast, the respirasome in different mammalian tissues does not contain COX7A2L and therefore lacks this tight interaction ^51^. Based on these structural data it was proposed that supercomplex CIII_2_-CIV is not a precursor of the respirasome ^51^. In contrast, a recent report proposes a role for COX7A2L in the formation of an alternate respirasome, suggested to be necessary for maintaining OXPHOS capacity upon metabolic switch to glycolysis ^52^. In the present study, we used the C57BL/6N mouse strain that is well documented to lack supercomplex CIII_2_-CIV ^9,47,48^. The homozygous *Uqcrc1* knock-in mice are thus deficient in both stable respirasomes and the CIII_2_-CIV supercomplex, which shows that CIII_2_ does not have to form any specific, tight interactions with other complexes to maintain OXPHOS activity.

The *Uqcrc1* knock-in mouse model we describe here has lost a stabilizing interaction between CI and CIII_2_, which otherwise locks them into a tightly defined superstructure as part of the respirasome. Upon disruption of this interaction CI and CIII_2_, which remain independently unaffected and fully functional, are free to take up alternate arrangements in the membrane. Supercomplexes are traditionally considered as spatially defined entities that remain associated when they are removed from the membrane environment. Our results argue that this type of stable respirasome arrangement is dispensable for bioenergetics and that respiratory chain complexes in proximity to each other in the membrane, but without having structurally defined relationships, can still sustain normal bioenergetics and physiology. Our results are also consistent with biochemical and structural studies showing that Q and Cyt *c* are not sequestered and restricted to exchange through structurally confined channels contained within individual respirasomes but are free to escape and exchange between them ^6,7,13,53^.

It is possible that respirasome formation ensures an even distribution of complexes throughout the membrane, keeping the diffusion distances and therefore timescales short through a proximity effect. Our results suggest that, if this is the case, the even distribution does not rely on tight, specific interactions, but may be promoted by the membrane topology and enzyme tessellation. Supercomplexes have also been proposed to prevent aggregation and nucleation of proteins in the very protein-rich inner mitochondrial membrane ^13,29,54^. This hypothesis is based on the idea that ordered protein-protein interactions prevent stronger deleterious interactions from occurring. A similar model has been proposed for the lens of the eye where ordered interactions between crystallin molecules prevent unwanted protein aggregation and cataract development ^55^.

It has been proposed that respirasomes may prevent electron leak and minimize ROS production at the level of the respiratory chain. One set of experiments favoring this model is based on measurements of ROS production at the level of CI after disrupting the CI-CIII_2_ interaction in bovine heart mitochondria or liposomes with the strong detergent DDM ^56^. Additional support for this model comes from a comparison of the levels of supercomplexes and ROS production in astrocytes and neurons, showing a positive correlation between the amount of free CI and ROS production ^57^. At variance with this proposed mechanism, structural data show that the flavin site of CI, one of two main sites for ROS production in the respiratory chain, is not protected within the respirasome structure but rather fully exposed to the mitochondrial matrix ^29^. In agreement with structural data, our experimental results do not provide evidence for increased ROS production when respirasomes are depleted.

Studies of human cell lines with mutations in mtDNA-encoded respiratory chain subunits have suggested that optimal maturation of CI occurs in the context of respiratory chain supercomplexes that serve as a platform for CI maturation ^58–60^. We show here that CI can be efficiently assembled despite very low levels of stable respirasomes, arguing that specific, stabilizing interactions between the different complexes may be dispensable for the efficacy of the CI assembly process, or, alternatively, that very low levels of respirasomes are sufficient to promote assembly of individual complexes.

Respirasomes have been observed in a range of eukaryotic species and we therefore expect that they provide a selective advantage, although our data argue that the mitochondrial bioenergetics in the mammalian membrane is not altered when they are destabilized. The *Uqcrc1* knock-in mouse model we describe here will be a valuable tool for understanding the role of respirasomes under various stress conditions, including different metabolic states, cancer, common age-associated degenerative diseases, and ageing.

### Limitations of study

The C57BL/6N mouse strain used for this study does not express the full-length COX7A2L (SCAFI) protein and therefore cannot form the supercomplex III_2_/IV as well as the recently reported proposed alternate respirasome. The role for COX7A2L in conjunction with the *Uqcrc1* knock-in allele needs to be further addressed in future studies.

Respirasomes may confer a selective advantage under certain stress conditions and further physiological assessment of *Uqcrc1* knock-in mice are therefore motivated.

The XL-MS approach has limitations, and the results should not be over-interpreted. The inner mitochondrial membrane contains ∼70% protein and the respiratory chain complexes are very abundant and will likely be close to each other regardless of if they form stable respirasomes or not. It is likely that many of the observed cross-links between respiratory chain complexes may be the result of transient interactions between the complexes or the consequence of the complex mitochondrial ultrastructure. Future studies will be needed to correlate the abundance of respirasomes (e.g., by cryo-electron tomography or focused ion beam scanning electron microscopy, FIB-SEM, of mitochondria) to provide a basis for a quantitative interpretation of the XL-MS data.

## ACKNOWLEDGEMENTS

N.G.L. was supported by the Swedish Research Council (2015-00418), Swedish Cancer Foundation, the Knut and Alice Wallenberg foundation (2016.0050 and 2019.0109), European Research Council (Advanced Grant 2016-741366), grants from the Swedish state under the agreement between the Swedish government and the county councils (SLL2018.0471) and the Max Planck Society. J.H. and I.C. were supported by the Medical Research Council (MC_U105663141 and MC_UU_00015/2). J.F.H. and A.J.R.H. acknowledge the Netherlands Organization for Scientific Research (NWO) for funding the Netherlands Proteomics Center through the X-omics Road Map program (project 184.034.019), and the Spinoza Grant SPI.2017.028. We would like to thank Petra Kirschner and Martin Purrio for expert technical assistance.

## AUTHOR CONTRIBUTIONS

N.G.L., J.H. and D.M. conceptualized research goals and experiments; D.M. planned, performed and analysed results from the majority of experiments; J.M. planned and performed RotaRod and open filed phenotyping experiments with help of R.F., A.J.R.H and J.F.H designed, performed and analysed cross-linking mass spectrometry experiments with help of J.M.; A.Mo. and T.M. planned, performed and analysed respiration, ATP and ROS production, of intact and permeabilized heart and liver mitochondria. I.C. planned, performed and analysed activity assays and BN-PAGE of mouse mitochondrial membranes. I.A. analyzed and interpreted complexome proteomics data and performed differential abundance analysis. X.L. performed complexome proteomics experiments and acquired mass spectrometry proteomics data. A.Me. performed, analysed and interpreted exercise performance related phenotyping experiments. N.G.L., J.H., D.M. and J.F.H wrote the manuscript, with input from all authors.

## MATERIAL AND METHODS

### Animals and housing

Knock-in and transgenic mice on a pure C57BL/6N background were housed in standard individually ventilated cages (45 × 29 × 12 cm) under a 12 h light/dark schedule (lights on 6 a.m. to 6 p.m.) in controlled environmental conditions of 22 ± 2 °C and 50 + 10% relative humidity. Normal chow diet and water were provided *ad libitum*. The study was approved by the Landesamt für Natur, Umwelt und Verbraucherschutz Nordrhein– Westfalen (reference numbers AZ.:84-02.04.2015.A103, AZ.: 84-02.05.50.15.004 and AZ.: 84-02.04. 2016.A420) and performed in accordance with the recommendations and guidelines of the Federation of European Laboratory Animal Science Associations (FELASA). Animal experiments in Sweden were approved by the animal welfare ethics committee and performed in compliance with national and European law.

### Generation of conditional knock-in mice

The targeting vectors for introducing mutations in *Ndufa11*, *Ndufb7* and *Uqcrc1* in embryonic stem cells were generated using BAC clones from the C57BL/6J RPCIB-731 BAC library and transfected into the Taconic Artemis C57BL/6N Tac ES cell line. Mice were generated by Taconic Artemis.

To establish a conditional knock-in allele for *Ndufa11,* a targeted allele was created to carry the wild-type exon 4 flanked by loxP sites and followed by a duplication of exon 4 with an Y110A coding mutation (TAT changed to GCA). The puromycin resistance cassette was introduced as a selection marker and removed by mating of *Ndufa11^+/loxP-pur^* mice with transgenic mice ubiquitously expressing *Flp*-recombinase. To obtain heart-and skeletal muscle-specific knock-in mice, *Ndufa11^loxP/loxP^* mice were crossed with transgenic mice expressing *cre*-recombinase under the control of the muscle creatinine kinase promoter (Ckmm*-cre*).

To establish a conditional knock-in allele for *Ndufb7*, a targeted allele was created where the wild-type exons 2 and 3 were followed by a duplicated region with exon 2 and a mutant version of exon 3 lacking last the last 60 nucleotides. The puromycin resistance cassette was removed by mating of *Ndufb7^+/loxP-pur^* mice with transgenic mice ubiquitously expressing *Flp*-recombinase. To obtain heart and skeletal muscle knockout mice *Ndufb7 ^loxP/loxP^* mice were crossed with Ckmm*-cre* transgenic mice.

To establish a conditional knock-in allele for *Uqcrc1*, the wild-type exons 7 and 13 were flanked by loxP sites and followed by a duplicated genomic region of exons 7-13 carrying nine nucleotide deletion (E258-D260) in exon 7. The puromycin resistance cassette was removed by mating of *Uqcrc1^+/loxP-pur^* mice with transgenic mice ubiquitously expressing *Flp*-recombinase. In a first set of experiments, heart-and skeletal muscle-specific knockout mice were generated by mating *Uqcrc1^loxp/loxP^* mice to Ckmm*-cre* transgenic mice. In subsequent studies, *Uqcrc1^+/loxP^* mice were mated with mice ubiquitously expressing *cre*-recombinase (β-actin-*cre*) to generate heterozygous knockout *Uqcrc1^+/DEL:E258-D260^* mice. Homozygous knockout *Uqcrc1^DEL:E258-D260/DEL:E258-D260^*mice were generated by intercrossing heterozygous knockout mice lacking the β-actin- *cre* transgene was removed by breeding.

### Isolation of mitochondria from mouse tissues

Mitochondria were isolated from mouse tissues using differential centrifugation. Freshly obtained tissues were cut, washed with ice cold PBS and homogenized in mitochondrial isolation buffer containing 320 mM sucrose, 10 mM Tris/HCl pH 7.4, and 1 mM EDTA and supplemented with 1× Complete protease inhibitor cocktail (Roche) by using a Teflon pestle (Schuett Biotec). After centrifugation at 1,000×*g* (swing-out rotor) for 10 min at 4 °C the supernatants were subsequently spun at 10,000×*g* for 10 min at 4 °C to isolate the mitochondria. Crude mitochondrial pellets were resuspended in suitable amount of mitochondrial isolation buffer.

For isolation of heart mitochondria, the following protocol was used. After removing all blood traces, freshly isolated heart tissue was cut into pieces on ice in a small volume of MIB (225 mM sucrose; 20 mM Tris/HCl pH 7.2; 1 mM EGTA). Heart pieces were collected and placed in a 2 ml Eppendorf tube containing 1.8 ml MIB and 200 µl of trypsin (2.5 %). After incubation for 10 minutes at 4 °C, 3 ml of MIB containing 0.04% trypsin inhibitor were added and homogenization (40 strokes, 450 rpm) was performed. The volume of homogenates was adjusted to 10 ml with MIB containing the trypsin inhibitor and samples were spun at 1,000×*g* for 10 min at 4 °C (swing out rotor). Next, the supernatants were collected and centrifuged at 10,000×*g* for 10 min at 4 °C using a fixed rotor centrifuge. The mitochondria-containing pellet was carefully resuspended in MIB buffer with 0.2% BSA and spun again 10,000×*g* for 10 min at 4 °C using a fixed rotor centrifuge. After washing the pellet with MIB and additional centrifugation (10,000×*g*, 10 min, 4 °C), mitochondrial pellets were resuspended in an appropriate volume of MIB.

Isolation of mitochondria from skeletal muscle tissue was performed as follows. All isolation steps were carried out on ice. Skeletal muscle tissue was minced into small pieces using scissors. Minced muscle was washed twice with ice-cold PBS/10mM EDTA by centrifugation (200×*g* 5 min, swing-out rotor) and subsequently resuspended in 5 ml of ice-cold PBS containing 10 mM EDTA and 0.05% trypsin for 30 min. Upon centrifugation at 200×*g* for 5 min (swing-out rotor) the pellet was resuspended in 10 ml IBm1 buffer (67 mM sucrose; 50 mM Tris/HCl pH 7.4; 50 mM KCl; 10 mM EDTA; 0.2% BSA) and homogenized using a Teflon pestle at 1300 rpm and 10-20 strokes. Upon adjusting the volume to 50 ml with IBm1 and centrifugation at 700×*g* for 10 min using a swing-out rotor the upper phase was transferred to a new tube and the whole procedure was repeated. Next, the supernatant was centrifuged at 8,000×*g* for 10 min (fixed rotor) and the pellet obtained resuspended in 10 ml of ice-cold IBm2 (250 mM sucrose; 3 mM EGTA, 10mM Tris/HCl pH 7.4). After a further centrifugation step of 8,000×*g* for 10 min (fixed rotor), the mitochondrial pellet was resuspended in an appropriate amount of IBm2. Protein concentrations were determined by the Bradford method using BSA as a standard.

### Preparation of mouse mitochondrial membranes

Membranes were prepared from heart mitochondria as described previously^63^. Briefly, for each membrane preparation, mitochondria from 2-3 mouse hearts were pooled and diluted to 5 mg protein ml^-1^ in 20 mM Tris/HCl (pH 7.55 at 20 °C), 1 mM EDTA, 10% glycerol, and ruptured using a Q700 Sonicator (Qsonica) with three 5 s bursts at 65% amplitude setting in 30 s intervals. Membranes were then pelleted by centrifugation (75,000x*g*, 1h) and resuspended in buffer.

### Activity assays of mouse mitochondrial membranes

All assays were conducted at 32 °C in 96-well plates using a Molecular Devices Spectramax 384 plus plate reader. Linear maximum rates were measured for all assays. For NADH:O_2_ oxidoreduction assays, membranes were diluted to 20 µg ml^-1^ in 10 mM Tris/H_2_SO_4_ (pH 7.5 at 32 °C) and 250 mM sucrose, supplemented with 3 µM horse heart Cyt *c* and 10 µg ml^-1^ alamethicin. Catalysis was initiated by addition of 200 µM NADH, and monitored at 340 and 380 nm (ε_340-380_ = 4.81 mM^-1^ cm^-1^). Inhibitor-insensitive rates (using 1 µM piericidin A) were subtracted from each measured rate. Succinate:O_2_ oxidoreduction was determined indirectly by monitoring NADPH (ε_340-380_ = 4.81 mM^-1^ cm^-1^) generated from a coupled enzyme assay in the presence of 5 mM succinate, 60 µg mL^-1^ FumC, 300 µg ml^-1^ MaeB, 2 mM MgSO_4_, 1 mM K_2_SO_4_ and 2 mM NADP^+ 64^. For measurement of superoxide (reactive oxygen species) production ^65^, membranes were diluted to 100 µg ml^-1^, supplemented with 10 µg ml^-1^ alamethicin, 10 units ml^-1^ superoxide dismutase (SOD) from bovine erythrocytes, 2 units ml^-1^ horseradish peroxidase (HRP), 10 µM Amplex Red and 1 µM piericidin A, and initiated using 30 µM NADH. H_2_O_2_ disproportionated from superoxide by SOD was quantified by monitoring the HRP-dependent oxidation of Amplex Red to resorufin at 557 and 620 nm (ε_557-620_ = 51.6 mM^-1^ cm^-1^). Catalase-insensitive rates (using 5,000 units ml^-1^ catalase from *Corynebacterium glutamicum*) were subtracted from each measured rate.

### Respiration on permeabilized mitochondria

Oxygen consumption of crude mitochondria permeabilized by cycles of freezing and thawing (5 cycles) was recorded as previously described^48^ using a high resolution O2K-oxygraph. Oxygen consumption was measured using 50 µg of heart mitochondria diluted in 2.1 ml of potassium phosphate buffer (50 mM, pH 7.2) and incubated with Cytochrome *c* from bovine heart (62.5 µg ml^-1^), NADH (1.2 mM) and/or succinate (10 mM).

### High-resolution oxygen consumption measurement

Oxygen consumption of intact crude mitochondria^18^ was measured at 37°C, using respectively 50 µg of heart mitochondria diluted in 2.1 ml of respiratory buffer (RB : 120 mM sucrose, 50 mM KCl, 20 mM Tris-HCl, 4 mM KH_2_PO_4_, 2 mM MgCl_2_, 1 mM EGTA, 0.25 mg/ml delipidated BSA; pH 7.2) in an Oxygraph-2k (Oroboros Instruments, Innsbruck, Austria). Oxygen consumption was measured by using different association of substrates: succinate (10 mM, pH 7.2), sodium pyruvate (10 mM, pH 7.2), L-glutamate (5 mM, pH 7.2), L-malate (5mM, pH 7.2). Oxygen consumption rate was measured under three different conditions: in the phosphorylating state with addition of ADP (5 mM, pH 7.2), in the non-phosphorylating state with addition of oligomycin (50 ng ml^-1^), and in the uncoupled state by successive addition of carbonyl cyanide m-chlorophenyl hydrazone (CCCP) up to 0.5 µM to reach the maximal respiration of the respiratory chain.

### ATP production measurement

ATP production rate and oxygen consumption of intact heart mitochondria (25µg/ml) was assessed in presence of ADP (5mM) and CI substrates: sodium pyruvate (10 mM; pH 7.2), L-glutamate (5 mM; pH 7.2), L-malate (5mM; pH 7.2) or CII substrates succinate (10 mM; pH 7.2) with the addition of CI inhibitor (rotenone, 30 nM). Mitochondria were incubated in RB (see “High-resolution oxygen consumption measurement”) in the presence of respiratory substrate as described above at 37°C. Aliquots were collected every 1 min and were precipitated with perchloric acid (PCA 7% without EDTA). After that, aliquots were centrifuged 20,000g for 10 min at 4 °C, and then were neutralized (pH 7.2) with 2 M KOH and 0.5 M MOPS. ATP content in these samples was determined with ATPlite 1step (PerkinElmer ^®^). The OXPHOS yield is defined by the ratio of ATP production rate per oxygen consumption rate, assessed in the same condition.

### H_2_O_2_ production measurement

H_2_O_2_ production of intact heart mitochondria (25µg/ml) were determined by monitoring the oxidation of the fluorogenic indicator Amplex red (Life Technologies) in the presence of horseradish peroxidase. The concentrations of horseradish peroxidase and Amplex red in the incubation medium were 12.5 U/ml and 25 nM, respectively. Fluorescence was recorded at the following wavelengths: excitation, 570 nm; emission, 582 nm with a spectrofluorimeter FP-8500 (Jasko). Mitochondria were incubated in the RB (see “High-resolution oxygen consumption measurement”) at 37°C. H_2_O_2_ production rate measurements were initiated by substrate addition and was measured for the three respiratory state previously described (see “High-resolution oxygen consumption measurement”) and for a fourth condition when the complex III is inhibited by antimycine A (100nM).

## BN-PAGE

For BN-PAGE, 75-100 μg of isolated mitochondria were lysed in 50 μl solubilization buffer (20 mM Tris pH 7.4; 0.1 mM EDTA; 50 mM NaCl; 10% [v/v] glycerol) containing 1-3% (w/v) digitonin (Calbiochem) or n-dodecyl β-D-maltoside (DDM, Glycon) and mixed with loading dye (5% [w/v] Coomassie Brilliant Blue G-250, 150 mM Bis-Tris, and 500 mM ε-amino-n-caproic acid [pH 7.0]). BN-PAGE samples were resolved on self-made 3%– 13% gels. BN-PAGE of mouse mitochondrial membranes was performed as follows. Mitochondrial membrane samples were resuspended to 5 mg mL^-1^ in 1.5 M aminocaproic acid, 50 mM Bis-Tris/HCl (pH 7.0) and solubilized on ice using 2% DDM or 1.67% digitonin for 20 mins. Protein concentrations were determined following centrifugation (48,000 *g*, 30 mins, 4 °C), and the supernatant mixed with NativePAGE sample buffer (Invitrogen) and G250 (to give 0.25x % detergent). Samples were centrifuged (14,000 *g*, 20 mins, 4 °C) to remove excess precipitated Coomassie, 10 µg (DDM-solubilized) or 15 µg (digitonin-solubilized) loaded in each lane of 3-12% Bis-Tris NativePAGE gels (Invitrogen), and run according to the manufacturer’s instructions. Proteins were visualized using Coomassie R250 staining, western blot analysis or *in gel* activity staining for complexes I, II and IV.

***In gel* catalytic activity assays**

For CI *in gel* activity ^48,66,67^, the BN-PAGE gel was incubated in 2 mM Tris/HCl pH 7.4, 0.1 mg ml^-1^ NADH (Roche) and 2.5 mg ml^-1^ iodonitrozolium for red staining or nitrotetrazolium blue for blue staining (Sigma) for about 10 minutes. CIV *in gel* activity was determined by incubating the BN-PAGE gels in 10 ml of 0.05 mM phosphate buffer pH 7.4, 25 mg 3.3´-diamidobenzidine tetrahydrochloride (DAB), 50 mg Cyt *c*, 3.75 g Sucrose and 1 mg Catalase for approximately 1h. The CII assay buffer contained 200 μl of sodium succinate (1 M), 8 μl of phenazine methosulfate (250 mM dissolved in DMSO), and 25 mg of NTB in 10 ml of 5 mM Tris/HCl, pH 7.4. Incubation of 10–30 min was required. The reactions were stopped using solution containing 50% methanol, 10% acetic acid for 30 min. All reactions were carried out at room temperature.

### Total heart proteomics

Five wild-type mouse hearts and five *Uqcrc1^DEL:E258-D260^* mouse hearts were used for the analysis. The frozen heart tissue was ground with a mortar and pestle under liquid nitrogen. A half full small spatula of the powder was transferred to a pre-cooled tube and an appropriate volume of guanidinium hydrochloride lysis buffer (6 M guanidinium chloride, 10 mM TCEP, 40 mM chloroacetamide and 100 mM Tris-HCl) was added. The tissues were lysed and in-solution digested with trypsin as previously described^68^ Tryptic peptides were desalted with C18 STAGE tips and eluted with 40% acetonitrile (ACN) /0.1% formic acid (FA). Four micro grams of the eluted peptides were dried and reconstituted in 9 µl of 0.1M TEAB. Labelling with Tandem Mass Tags (TMT, TMT10plex, Thermo Fisher Scientific cat. No 90110) was carried out according to manufacturer’s instruction with the following changes: 0.8 mg of TMT reagent was re-suspended with 70 µl of anhydrous ACN. Seven micro litters of TMT reagent in ACN was added to 9 µl of peptide resuspended in 0.1M TEAB. The final ACN concentration was 43.75% and the ratio of peptides to TMT reagent was 1:20. After 60 min of incubation, the reaction was quenched with 2 µl of 5% hydroxylamine. Labelled peptides were pooled, dried, re-suspended in 200 µl of 0.1% formic acid (FA), split into two equal parts, and desalted using home-made STAGE tips^69^. One of the two parts was fractionated on a 1 mm x 150 mm ACQUITY column, packed with 130 Å, 1.7 µm C18 particles (Waters cat. no SKU: 186006935), using an Ultimate 3000 UHPLC (Thermo Fisher Scientific). Peptides were separated using a 96 min segmented gradient from 1% to 50% buffer B for 85 min and from 50% to 95% buffer B for 11 min, at a flow of 30 µl/min with a; buffer A was 5% ACN, 10mM ammonium bicarbonate (ABC), buffer B was 80% ACN, 10mM ABC. Fractions were collected every three minutes, and fractions were pooled in two passes (1 + 17, 2 + 18 … etc.) and dried in a vacuum centrifuge (Eppendorf). Dried fractions were re-suspended in 0.1% formic acid (FA) and separated on a 50 cm, 75 µm Acclaim PepMap column (Product No. 164942 Thermo Fisher Scientific) using an EASY-nLC1200 (Thermo Fisher Scientific). The analytical column was operated at 50°C. The separation was performed using a using a 90 min linear gradient from 6% to 31% buffer; buffer A was 0.1% FA, buffer B was 0.1% FA, 80% ACN. Eluting peptides were analysed on an Orbitrap Lumos Tribrid mass spectrometer (Thermo Fisher Scientific) equipped with a FAIMS device (Thermo Fisher Scientific). The FAIMS device was operated at two compensation voltages: −50 V and −70 V. Mass spectrometric data were acquired in a data-dependent manner with a top speed method. For MS1, the mass range was set to 350−1500 m/z and resolution to 60K. Maximum injection time was 50 ms and the AGC target to 4e5. Peptides were fragmented using collision-induced dissociation; collision energy was to 35%. Peptide fragment MS2 spectra were acquired in the ion trap with a maximum injection time of 50 ms and “Turbo” scan rate, using an AGC target of 1e4. The ten most abundant peaks were subjected to synchronous precursor selection and fragmented using higher-energy collisional dissociation; collusion energy was set to 65%. The resulting MS3 spectra were acquired in the Orbitrap at a resolution of 50K. Raw files were split based on the FAIMS compensation voltage using FreeStyle (Thermo Fisher Scientific).

Raw data were analysed using MaxQuant version 1.6.10.43 as described below except that quantification was set to “Reporter ion MS3”. The isotope purity correction factors, provided by the manufacturer, were included in the analysis. Differential expression analysis was performed using limma^70^.

### Complexome profiling

Each BN-PAGE gel lane was cut into 24 equal-sized bands with a sharp scalpel and each band was chopped into 1 mm^3^ pieces. The pieces from each band were placed into a well of an OASIS® HLB µElution Plate (Waters, Milford, MA, USA). The gel bands were de-stained, reduced, alkylated, tryptic digested and desalted in the OASIS® HLB µElution Plate with the help of a Positive Pressure-96 Processor (Waters, Milford, MA, USA) ^71^. The cleaned peptides were re-suspended with 0.1% formic acid analysed by mass spectrometry. Peptides were analysed using an Orbitrap Fusion mass spectrometer (Thermo Fisher Scientific) with a nano-electrospray ion source, coupled with an EASY-nLC 1000 (Thermo Fisher Scientific) UHPLC. A 25 cm long reversed-phase C18 column with 75 μm inner diameter (PicoFrit, LC Packings) was used for separating peptides. The flowing rate was 250nl min^-1^. The LC runs lasted 31 min with a concentration of 6% solvent B (0.1% formic acid in 80% acetonitrile) increasing to 31% over 27 min and further to 44% over 4 min. The column was subsequently washed and re-equilibrated. MS spectra were acquired in a data-dependent manner with a top speed method. For MS, the mass range was set to 350−1500m/z and resolution to 60K at 200m/z. The AGC target of MS was set to 1e6, and the maximum injection time was 50 ms. Peptides were fragmented with HCD with collision energy of 27% and resolution of 30K. The AGC target was set to 2e5 and the maximum injection time was 80 ms.

The raw data were analysed with MaxQuant version 1.6.0.13^72^ using the integrated Andromeda search engine^73^eptide fragmentation spectra were searched against the canonical sequences of the mouse reference proteome (UniProt proteome ID UP000000589, downloaded September 2018). Methionine oxidation and protein N-terminal acetylation were set as variable modifications; cysteine carbamidomethylation was set as fixed modification. The digestion parameters were set to “specific” and “Trypsin/P,” The minimum number of peptides and razor peptides for protein identification was 1; the minimum number of unique peptides was 0. Protein identification was performed at a peptide spectrum matches and protein false discovery rate of 0.01. The “second peptide” option was on. The “Match between runs” option was enabled to transfer identifications within the gel slices of a single BN lane only. Each BN gel lane slice was designated as a separate experiment. Protein intensities were normalized to the highest value per BN lane. Exploratory data analysis and visualization was done using ggplot ^74^ in R^75^.

MaxQuant’s MaxLFQ algorithm ^31^ was used for label-free quantification (LFQ) of whole protein complexes; an LFQ ratio count of 2 was selected. Differences in abundance, per BN-PAGE slice, of subunits or whole protein complexes were calculated using limma^70^. The whole complex intensity was calculated from the sum of intensity of all corresponding subunits. Statistical analysis was performed using limma’s moderated t-test.

For the quantification of individual proteins, i.e., respiratory chain assembly factors, only proteins with at least three reported intensity values were used for analysis. Missing values were imputed from a normal distribution with a downshift of 1.8 and a standard deviation of 0.3. Testing for significant differences in abundance between the corresponding BN-PAGE slice was done using Limma’s moderated t-test.

### Western blot analysis

Proteins were separated by SDS–PAGE or BN–PAGE and then transferred to polyvinylidene difluoride (PVDF) membranes (Milipore). Immunodetection was performed according to the standard techniques using enhanced chemiluminescence (Immun-Star HRP Luminol/Enhancer Bio Rad) or fluorescence. To perform the fluorescent detection of complex I and III, PVDF membranes were blocked using the Rockland blocking buffer (MB-070). After incubation with the primary antibody, an Alexa Fluor 680 goat anti-rabbit or an IRdye800 anti-mouse secondary antibody was used. The detection was performed using the Li-COR Odyssey system. Abcam antibodies specific for OXPHOS (ab110413), UQCRC1 (ab 118687), UQCRFS1 (ab14746), VDAC (ab14734) were used. Antibodies against SDHA (459200) and NDUFB7 (PA5-51743 Thermo Fisher) were obtained from Invitrogen. NDUFV1 antibody was purchased from Proteintech (11238-1- AP).

### Cross-linking of mouse heart mitochondria

For XL-MS, wild-type and *Uqcrc1* knock-in mitochondria purified from three biological replicates were treated with an optimal cross-linker concentration of 0.5 mM for 45 minutes at 15 °C (Figure S7A) Cross-linking experiments showed high reproducibility across replicates and conditions (Figure S7B). The cross-link reaction was quenched for 30 min at 25 °C by the addition of 50 mM Tris (1 M Tris buffer, pH 8.5). Subsequently, cross-linked mitochondria were pelleted at 11,000 g at 4 °C for 10 min and re-suspended in mitochondrial lysis buffer (100 mM Tris pH 8.5, 7 M Urea, 1 % Triton-X-100, 5 mM TCEP, 30 mM CAA, 2.5 mM Mg^2+^, proteinase inhibitor cocktail). Mitochondria were solubilized for 30 min on ice. Next, proteins in the soluble fraction were precipitated using Methanol/Chloroform precipitation as previously described^76^. The dried protein pellet was then re-suspended in digestion buffer (100 mM Tris-HCl pH 8.5, 1% sodium deoxycholate, 5 mM TCEP, and 30 mM CAA). Protein digestion was performed overnight at 37 °C using Trypsin (1:25 ratio weight/weight) and Lys-C (1:100 ratio weight/weight), respectively. Finally, peptides were de-salted solid-phase extraction C18 columns (Sep-Pak, Waters) and fractionated into 22 fractions using an Agilent 1200 HPLC pump system (Agilent) coupled to a strong cation exchange (SCX) separation column (Luna SCX 5 µm to 100 Å particles, 50 × 2 mm, Phenomenex).

### Cross-linking mass spectrometry and data analysis

The 22 SCX fractions of DSSO were analyzed using an Ultimate3000 (Thermo Fisher Scientific) and 50-cm analytical column packed with C18 beads (Dr Maisch Reprosil C18, 3 µm) heated at 45°C, connected to an Orbitrap Fusion Lumos. Peptides were eluted at 300 nL/min with a 95-minute gradient from 9% to 40% of buffer B (80% Acetonitrile and 20% water with 0.1% formic acid). To identify cross-linked peptides, MS1 scans with the Orbitrap resolution set to 120,000 from 350 to 1400 m/z were performed (normalized AGC target set to 250 % and maximum injection time set to auto). Ions with a charge from +3 to +8 were fragmented with stepped HCD (21 %, 27 %, 33 %) and further analysed (MS2) in the Orbitrap (30,000 resolution, normalized AGC target set to 200 % and maximum injection time set to auto) for detection of DSSO signature peaks (difference in mass of 37.972 Da). The four ions with this specific difference were further sequenced following MS3 CID - Ion Trap scans (collision energy 35 %, normalized AGC target set to 200 % and maximum injection time set to auto). The raw files corresponding to the 22 DSSO fractions were analysed with Proteome Discoverer software suite version 2.5 (Thermo Fisher Scientific) with the incorporated XlinkX node for analysis of cross-linked peptides as reported by Klykov et al ^77^. Data were searched against a FASTA file containing proteins, which were previously identified following a classical bottom-up workflow. Were applicable, mitochondrial target peptides were removed from respective protein sequences. For the XlinkX search, fully tryptic digestion with three maximum missed cleavages, 10 ppm error for MS1, 20 ppm for MS2 and 0.5 Da for MS3 in Ion Trap was set as search parameters. Carbamidomethyl (C) was set as static modification while Oxidation (Met) and acetylation (protein n-terminus) were set as dynamic modification. Cross-linked peptides were accepted with a minimum XlinkX score of 40 and maximum FDR (controlled at PSM level for cross-linked spectrum matches) rate set to 5%.

### Identification of respirasome complex inter links and MS1 mass tags

Identified inter cross-links between the individual members of the respirasome complex were mapped onto a modified respirasome complex structure using ChimeraX. Briefly, structures for mouse complex I, complex III and complex IV (CI = PDB 6ZR2, CIII = PDB 7O3H), CIV = PDB 7O3E) were structurally aligned based on the previously resolved respirasome structure of Sus scrofa (5GUP). Additionally, unresolved, cross-linked N-termini are modelled using Alphafold^78^. Identified cross-links were classified as respirasome structure-compatible and in-compatible using a distance threshold (Cα-Cα distance <= 30 Å).

To additionally quantify respirasome cross-linking frequencies in wild-type and knock-in samples, an in-depth analysis of the recorded ion mass spectra (MS1 analysis) of identified cross-linked peptide precursors for respirasome structure-compatible and respirasome structure-incompatible crosslinks was performed. Briefly, to identify DSSO linked peptides, highly abundant ions detected in the MS1 scan are subsequently selected and fragmented in two consecutive scans (MS2 and MS3). As cross-linked peptide ions have very low abundance, only a very small fraction is fragmented and subsequently identified. Matching masses of identified cross-link peptides against all recorded masses (MS1 mass matching) allows the identification of non-fragmented ions, thereby providing a more complete estimate for the frequency of cross-linked peptides in the sample. Recorded raw files for wild-type and Uqcrc1 knock-in samples were de-convoluted with MaxQuant ^72^ to obtain two separate lists with all detected LC-MS features (allPeptides.txt). Respective lists were subsequently filtered for peptides with charge states above two. Next, deprotonated masses of respirasome structure-compatible and respirasome structure-incompatible cross-linked peptides were matched against filtered LC-MS features for either wild-type or knock-in samples using a custom python script. Peptides from the filtered feature lists with a resulting mass tolerance of +/- 0.03 Da were considered a mass match for an identified respirasome structure-compatible cross-linked peptide.

### Quantification of cross-linked peptides

Cross-linked peptide pairs were quantified by combining the MS1 precursor intensity and with spectral counts similar the workflow described by Chen *et al.* ^79^. Briefly, identified cross-linked peptide precursor intensities were first median normalized (considering the median intensity for each replicate and the global median intensity) and subsequently summed for respective cross-links. Cross-links for which no precursor abundance could be determined were disregarded.

### Polymerase chain reaction

For verification of nine nucleotide deletion (E258-D260) in *Uqcrc1*, we used PCR on genomic DNA isolated from heart of homozygous *Uqcrc1* knock-in and wild-type mice, with the following primers Uqcrc1_fw (TGGAACACCAGCAGCTGC) and Uqcrc1_rev (CACACCTCACTGCCAGTG). The products were analysed on 2,5% agarose gels.

### Body composition measurements

Body fat and lean mass content were measured *in vivo* by nuclear magnetic resonance using the minispec LF50H (Bruker Optics).

### Treadmill and running wheels exercise experiments

Mice were placed on a treadmill (TSE Systems) equipped with an electroshock grid for a 5-minute habituation period. After a 10-minute warm-up phase at 0.1 m/sec, the speed of the belt was continuously increased by 0.02 m/sec per minute until the mice were exhausted. Exhaustion was defined by three consecutive shocks (0.3 mA). For the running wheel experiment voluntary running of individually housed mice was recorded for 14 days using wireless running wheels (Med Associates Inc.)

### Rotarod and open field assays

All animals were allowed to acclimatize to the behaviour laboratory for 30-60 min prior to testing/training. Each test/training session occurred at the same time during the day (9 am-5 pm) with small variations. For the Rotarod (Ugo Basile), mice were trained twice the day before the actual assessment. During each training session mice spent 1.5 min on the rod rotating at the constant speed of 4 rpm. On the assessment day mice were put on the accelerating rod (4-40 rpm) and their latency to fall off the rod was measured. The maximum rod speed of 40 rpm was reached in 5 min, the maximum time mice were allowed to spend on the rod. Each animal had 5 trials with approximately 15 min break in between. For the openfield test, each animal was placed in the arena with the walls and allowed to freely explore the field for 60 min. The openfield test was performed using ActiMot detection system (TSE systems).

### Metabolic cages

Indirect calorimetry, food and water intake were measured for singly housed mice in metabolic cages (PhenoMaster, TSE Systems) under resting conditions. Prior to the experiments, mice were acclimatized to the housing conditions for 3 to 4 days. Afterwards mice were transferred to the metabolic cages and data were collected for 48 hours after at least 1 day of acclimatization in the metabolic cages. Locomotor activity was simultaneously recorded by the interruption of light beams.

### Data Availability

The mass spectrometry proteomics data and data analysis code have been deposited to the ProteomeXchange Consortium via the PRIDE partner repository with the dataset identifier PXD023846; Username, Password: nIuEY87x All cross linking mass spectrometry related data are available via the PRIDE partner repository with the dataset identifier PXD031345 (**Username:****Password:** LX46KYFd).

## SUPPLEMENTAL FIGURE LEGENDS

**Figure S1. Genomic targeting for generation of knock-in mice.** Targeting strategies for the conditional knock-in of the *Ndufa11* (A), *Ndufb7* (B) and *Uqcrc1* (C) genes. Red arrowheads represent *loxP* sites. Green arrowheads represent *frt* sites and blue arrowheads represent F3 sites that are used to remove selection markers by homologous recombination.

**Figure S2. The effect of different knock-in mutations on the mitochondrial OXPHOS supercomplexes.** (A) BN-PAGE analysis followed by CI *in gel* activity assay of wild-type and *Ndufa11* knock-in heart mitochondria using DDM or various concentrations of digitonin. (B) CI *in gel* activity staining of wild-type and *Ndufb7* knock-in heart mitochondria subjected to increasing digitonin concentrations. (C) Truncated version of the NDUFB7 protein visualized by SDS-PAGE and western blot. Antibodies used for detection are indicated. (D) Verification of the *Uqcrc1* knock-in mutations by PCR amplification of genomic DNA demonstrating the nine nucleotides deletion. PCR was performed on DNA isolated from heart tissue of wild-type and *Uqcrc1* knock-in mice. (E) Western blot analysis of steady-state levels of OXPHOS proteins from mitochondria isolated from denoted tissues of control and *Uqcrc1* knock-in mice. (F) Lack of changes in steady state levels of OXPHOS subunits in *Uqcrc1^DEL:E258-D260^* hearts; n=5. None of the subunits test significant for differences in abundance. The error bars represent the 95% confidence interval for the log2 fold change (logFC) value.

**Figure S3. Heat-map representing complexome profiling of OXPHOS protein distribution and abundance in wild-type and *Uqcrc1* knock-in mitochondria.** The complexome analysis is based on three replicates per genotype. Control complexes represent known mitochondrial protein complexes containing proteins unrelated to OXPHOS, e.g., VDAC, prohibitins and DNAJC15.

**Figure S4. Boxplot and volcano plot representation of complexome analysis of BN-PAGE area containing novel, *Uqcrc1* knock-in specific, complex.** The boxplots show the distribution of CI-CV subunits along the BN-PAGE gel area containing the novel aberrant complex seen in Uqcrc1 knock-in versus wild-type mitochondria. The volcano plots show all significantly enriched proteins along analysed fractions. Fraction number 3 is enriched for the novel aberrant complex. Fractions 1 and 2 contain higher molecular weight and fraction 4 and 5 lower molecular weight fractions surrounding fraction 3. (A) Heart, (B) Skeletal muscle, (C) Kidney.

**Figure S5. Bioenergetic properties of mitochondria are not compromised by the homozygous *Uqcrc1* knock-in mutation.** (A) Coomassie staining and *in gel* activity on isolated mitochondrial membranes from wild-type and *Uqcrc1* knock-in mice upon DDM (left) and digitonin (right) solubilization and BN-PAGE analyses. (B) H_2_O_2_ production rates in heart mitochondria of wild-type and *Uqcrc1* knock-in mice with CI and CII substrates. Statistical analyses were performed using one-way ANOVA. Error bars indicate mean ± SEM. (C) OXPHOS yield (JATP/JO_2_) was measured in intact heart mitochondria isolated from wild-type and *Uqcrc1* knock-in mice (n=9). Statistical analyses were performed using one-way ANOVA. Error bars indicate mean ± SEM. Abbreviations: PGM, pyruvate, glutamate and malate; Succ, succinate; rot, rotenone.

**Figure S6. Bioenergetic properties of mitochondria are not compromised by the homozygous *Uqcrc1* knock-in mutation.** (A) The maximal oxygen consumption rate assessed in permeabilized liver mitochondria from control (white bars) and *Uqcrc1* KI (grey bars) mice in the presence of non-limiting concentrations of substrates for complex I (I-III-IV), complex II (II-III-IV) or both (I-II-III-IV). n = 4; error bars indicate mean ± SEM. Statistical test: one-way ANOVA.

(B-C) Oxygen consumption of liver mitochondria from control (white bars) and *Uqcrc1* KI (grey bars) mouse strains at 22 weeks of age. n = 9; error bars indicate mean ±SEM. Statistical test: one-way ANOVA. Abbreviations: PGM, pyruvate, glutamate, and malate; Succ, succinate; rot, rotenone.

(D-E) Respiratory chain enzyme activity (CI, CII, CI-CIII, CII-CIV, CIV) measurements in isolated mitochondria from control (white bars) *Uqcrc1* KI (grey bars) mice for mitochondria isolated from heart (D) and liver (E). n = 9; Error bars indicate mean ± SEM. Statistical test: one-way ANOVA.

**Figure S7. Exercise and locomotion in *Uqcrc1* knock-in mice are not affected by very low levels of stable respirasomes.** (A) Schematic overview of the phenotyping experiment. Body weights (B) and fat content (C) of wild-type and *Uqcrc1* knock-in mice of both sexes before and after exercise (day 1 and day 14 of the experiment). The values represent mean values from two independent experiments with total number of mice n = 12 (male) and n = 8 (female). Error bars indicate mean ± SEM. (D) Treadmill distances run by wild-type and *Uqcrc1* knock-in mice of both sexes at three indicated time points. (E) Total distance of voluntary running on running wheels situated in the cage during the period of 15 days. The values in (D) and (E) are mean averages from three independent experiments with total number of mice n = 16 for males and n = 8 for females. Statistical analyses were performed using Welch’s t-test. Error bars indicate mean ± SEM.

**Figure S8. Metabolic cages** Water intake (A, B), food intake (C, D), respiratory exchange ratio (RER) (E, F), physical activity (G, H), oxygen consumption (VO_2_) (I, J) and CO_2_ production (VCO_2_) (K, L) of wild type and *Uqcrc1* knock-in mice of both sexes for 48 hours. Grey boxes indicate mouse housing in dark. For males n=9 per genotype; for females, n of wild type=6, n of *Uqcrc1* knock-in= 8. Statistical analyses were performed using Welch’s t-test. Error bars indicate mean ± SEM.

**Figure S9. Overview of cross-linking of wild-type and *Uqcrc1* knock-in mitochondria** (A) Coomassie-stained gel showing optimization of the conditions used for cross-linking experiment on intact mitochondria. (B) Count of cross-links and underlying spectral matches (CSM) identified for wild-type (blue) and Uqcrc1 knock-in (orange) replicates. The number of cross-links and CSMs with an identified precursor abundance (determining whether an observed cross-link can be quantified) is indicated in brackets. (C) Venn diagrams showing the reproducibility of cross-linking results between the three replicates and conditions (wild-type versus Uqcrc1 knock-in). (D) Count of cross-links observed between and within subunits (intra respirasome cross-links) of either complex I (light blue), complex III (dark blue) or complex IV (light green) in wild-type and *Uqcrc1* knock-in replicates. For validation, cross-links (black lines) were mapped on the respective mouse complex structures (PDB: 6ZR2, 7O3H, 7O3E), with cross-linked, previously unresolved N-termini being modelled using Alphafold^78^. Respective distances are shown in the boxplot as grey dots, with outliers marked with a dark grey diamond and the median cross-link distance indicated as grey line and on top of each plot. (E) Count of all recorded MS1 features with a charge state > 2 for wild-type (blue) and *Uqcrc1* knock-in (orange). For wild-type and *Uqcrc1* knock-in, MS1 features were combined from spectra across the three replicates. (F) Cross-link precursor abundance proteome wide and intra respirasome cross-links are shown as box plot for wild-type (blue) and Uqcrc1 knock-in (orange) replicates. For clarity of the box plot, outliers are not visualized.

## REFERENCES

1. Schägger, H., and Pfeiffer, K. (2000). Supercomplexes in the respiratory chains of yeast and mammalian mitochondria. EMBO J 19, 1777–1783. 10.1093/emboj/19.8.1777.

2. Chavez, J.D., Lee, C.F., Caudal, A., Keller, A., Tian, R., and Bruce, J.E. (2017). Chemical Crosslinking Mass Spectrometry Analysis of Protein Conformations and Supercomplexes in Heart Tissue. Cell Systems, 1–12. 10.1016/j.cels.2017.10.017.

3. Liu, F., Lössl, P., Rabbitts, B.M., Balaban, R.S., and Heck, A.J.R. (2018). The interactome of intact mitochondria by cross-linking mass spectrometry provides evidence for coexisting respiratory supercomplexes*. Mol Cell Proteomics 17, 216–232. 10.1074/mcp.ra117.000470.

4. Rieger, B., Shalaeva, D.N., Söhnel, A.-C., Kohl, W., Duwe, P., Mulkidjanian, A.Y., and Busch, K.B. (2017). Lifetime imaging of GFP at CoxVIIIa reports respiratory supercomplex assembly in live cells. Sci. Rep. 7, 46055. 10.1038/srep46055.

5. Davies, K.M., Blum, T.B., and Kühlbrandt, W. (2018). Conserved in situ arrangement of complex I and III2 in mitochondrial respiratory chain supercomplexes of mammals, yeast, and plants. Proc. Natl. Acad. Sci. U.S.A. 115, 3024–3029. 10.1073/pnas.1720702115.

6. Gu, J., Wu, M., Guo, R., Yan, K., Lei, J., Gao, N., and Yang, M. (2016). The architecture of the mammalian respirasome. Nature 537, 639–643. 10.1038/nature19359.

7. Letts, J.A., Fiedorczuk, K., and Sazanov, L.A. (2016). The architecture of respiratory supercomplexes. Nature 537, 644–648. 10.1038/nature19774.

8. Kobayashi, A., Azuma, K., Takeiwa, T., Kitami, T., Horie, K., Ikeda, K., and Inoue, S. (2023). A FRET-based respirasome assembly screen identifies spleen tyrosine kinase as a target to improve muscle mitochondrial respiration and exercise performance in mice. Nat Commun 14, 312. 10.1038/s41467-023-35865-x.

9. Lapuente-Brun, E., Moreno-Loshuertos, R., Acin-Perez, R., Latorre-Pellicer, A., Colas, C., Balsa, E., Perales-Clemente, E., Quiros, P.M., Calvo, E., Rodríguez-Hernández, M.A., et al. (2013). Supercomplex Assembly Determines Electron Flux in the Mitochondrial Electron Transport Chain. Science 340, 1567–1570. 10.1126/science.1230381.

10. Balsa, E., Soustek, M.S., Thomas, A., Cogliati, S., García-Poyatos, C., Martín-García, E., Jedrychowski, M., Gygi, S.P., Enriquez, J.A., and Puigserver, P. (2019). ER and Nutrient Stress Promote Assembly of Respiratory Chain Supercomplexes through the PERK-eIF2α Axis. Mol Cell 74, 877–890.e6. 10.1016/j.molcel.2019.03.031.

11. Acín-Pérez, R., Fernández-Silva, P., Peleato, M.L., Pérez-Martos, A., and Enríquez, J.A. (2008). Respiratory active mitochondrial supercomplexes. Mol. Cell 32, 529–539. 10.1016/j.molcel.2008.10.021.

12. Sousa, J.S., Mills, D.J., Vonck, J., and Kühlbrandt, W. (2016). Functional asymmetry and electron flow in the bovine respirasome. eLife 5, 805. 10.7554/elife.21290.

13. Blaza, J.N., Serreli, R., Jones, A.J.Y., Mohammed, K., and Hirst, J. (2014). Kinetic evidence against partitioning of the ubiquinone pool and the catalytic relevance of respiratory-chain supercomplexes. Proc. Natl. Acad. Sci. U.S.A. 111, 15735–15740. 10.1073/pnas.1413855111.

14. Trouillard, M., Meunier, B., and Rappaport, F. (2011). Questioning the functional relevance of mitochondrial supercomplexes by time-resolved analysis of the respiratory chain. Proc. Natl. Acad. Sci. U.S.A. 108, E1027–34. 10.1073/pnas.1109510108.

15. Vercellino, I., and Sazanov, L.A. (2021). The assembly, regulation and function of the mitochondrial respiratory chain. Nat Rev Mol Cell Bio, 1–21. 10.1038/s41580-021-00415-0.

16. Fedor, J.G., and Hirst, J. (2018). Mitochondrial Supercomplexes Do Not Enhance Catalysis by Quinone Channeling. Cell Metab 28, 525–531.e4. 10.1016/j.cmet.2018.05.024.

17. Moe, A., Trani, J.D., Rubinstein, J.L., and Brzezinski, P. (2021). Cryo-EM structure and kinetics reveal electron transfer by 2D diffusion of cytochrome c in the yeast III-IV respiratory supercomplex. Proc National Acad Sci 118, e2021157118. 10.1073/pnas.2021157118.

18. Molinié, T., Cougouilles, E., David, C., Cahoreau, E., Portais, J.-C., and Mourier, A. (2022). MDH2 produced OAA is a metabolic switch rewiring the fuelling of respiratory chain and TCA cycle. Biochimica Et Biophysica Acta Bba - Bioenergetics 1863, 148532. 10.1016/j.bbabio.2022.148532.

19. Novack, G.V., Galeano, P., Castaño, E.M., and Morelli, L. (2020). Mitochondrial Supercomplexes: Physiological Organization and Dysregulation in Age-Related Neurodegenerative Disorders. Front Endocrinol 11, 600. 10.3389/fendo.2020.00600.

20. Champagne, D.P., Hatle, K.M., Fortner, K.A., D’Alessandro, A., Thornton, T.M., Yang, R., Torralba, D., Tomás-Cortázar, J., Jun, Y.W., Ahn, K.H., et al. (2016). Fine-Tuning of CD8(+) T Cell Mitochondrial Metabolism by the Respiratory Chain Repressor MCJ Dictates Protection to Influenza Virus. Immunity 44, 1299–1311. 10.1016/j.immuni.2016.02.018.

21. Rosca, M., Minkler, P., and Hoppel, C.L. (2011). Cardiac mitochondria in heart failure: normal cardiolipin profile and increased threonine phosphorylation of complex IV. Biochim. Biophys. Acta 1807, 1373–1382. 10.1016/j.bbabio.2011.02.003.

22. McKenzie, M., Lazarou, M., Thorburn, D.R., and Ryan, M.T. (2006). Mitochondrial respiratory chain supercomplexes are destabilized in Barth Syndrome patients. J. Mol. Biol. 361, 462–469. 10.1016/j.jmb.2006.06.057.

23. Frenzel, M., Rommelspacher, H., Sugawa, M.D., and Dencher, N.A. (2010). Ageing alters the supramolecular architecture of OxPhos complexes in rat brain cortex. Exp. Gerontol. 45, 563–572. 10.1016/j.exger.2010.02.003.

24. Ikeda, K., Horie-Inoue, K., Suzuki, T., Hobo, R., Nakasato, N., Takeda, S., and Inoue, S. (2019). Mitochondrial supercomplex assembly promotes breast and endometrial tumorigenesis by metabolic alterations and enhanced hypoxia tolerance. Nat Commun 10, 4108. 10.1038/s41467-019-12124-6.

25. Rohlenova, K., Sachaphibulkij, K., Stursa, J., Bezawork-Geleta, A., Blecha, J., Endaya, B., Werner, L., Cerny, J., Zobalova, R., Goodwin, J., et al. (2017). Selective Disruption of Respiratory Supercomplexes as a New Strategy to Suppress Her2high Breast Cancer. Antioxid Redox Sign 26, 84–103. 10.1089/ars.2016.6677.

26. Hollinshead, K.E.R., Parker, S.J., Eapen, V.V., Encarnacion-Rosado, J., Sohn, A., Oncu, T., Cammer, M., Mancias, J.D., and Kimmelman, A.C. (2020). Respiratory Supercomplexes Promote Mitochondrial Efficiency and Growth in Severely Hypoxic Pancreatic Cancer. Cell Reports 33, 108231. 10.1016/j.celrep.2020.108231.

27. Huertas, J.R., Fazazi, S.A., Hidalgo-Gutierrez, A., López, L.C., and Casuso, R.A. (2017). Antioxidant effect of exercise: Exploring the role of the mitochondrial complex I superassembly. Redox Biol 13, 477–481. 10.1016/j.redox.2017.07.009.

28. Greggio, C., Jha, P., Kulkarni, S.S., Lagarrigue, S., Broskey, N.T., Boutant, M., Wang, X., Alonso, S.C., Ofori, E., Auwerx, J., et al. (2016). Enhanced Respiratory Chain Supercomplex Formation in Response to Exercise in Human Skeletal Muscle. Cell metabolism 25, 1–12. 10.1016/j.cmet.2016.11.004.

29. Milenkovic, D., Blaza, J.N., Larsson, N.-G., and Hirst, J. (2017). The Enigma of the Respiratory Chain Supercomplex. Cell metabolism 25, 765–776. 10.1016/j.cmet.2017.03.009.

30. Lobo-Jarne, T., and Ugalde, C. (2017). Respiratory chain supercomplexes: Structures, function and biogenesis. Semin. Cell Dev. Biol. 10.1016/j.semcdb.2017.07.021.

31. Cox, J., Hein, M.Y., Luber, C.A., Paron, I., Nagaraj, N., and Mann, M. (2014). Accurate Proteome-wide Label-free Quantification by Delayed Normalization and Maximal Peptide Ratio Extraction, Termed MaxLFQ*. Mol Cell Proteomics 13, 2513–2526. 10.1074/mcp.m113.031591.

32. Schägger, H. (2001). Respiratory chain supercomplexes. IUBMB Life 52, 119–128. 10.1080/15216540152845911.

33. Wu, M., Gu, J., Guo, R., Huang, Y., and Yang, M. (2016). Structure of Mammalian Respiratory Supercomplex I1III2IV1. Cell 167, 1598–1609.e10. 10.1016/j.cell.2016.11.012.

34. Hevler, J.F., Lukassen, M.V., Cabrera-Orefice, A., Arnold, S., Pronker, M.F., Franc, V., and Heck, A.J.R. (2021). Selective cross-linking of coinciding protein assemblies by in-gel cross-linking mass spectrometry. Embo J 40, e106174. 10.15252/embj.2020106174.

35. Kao, A., Chiu, C., Vellucci, D., Yang, Y., Patel, V.R., Guan, S., Randall, A., Baldi, P., Rychnovsky, S.D., and Huang, L. (2011). Development of a Novel Cross-linking Strategy for Fast and Accurate Identification of Cross-linked Peptides of Protein Complexes*. Mol Cell Proteomics 10, M110.002170. 10.1074/mcp.m110.002212.

36. Bomba-Warczak, E., Edassery, S.L., Hark, T.J., and Savas, J.N. (2021). Long-lived mitochondrial cristae proteins in mouse heart and brain. J Cell Biol 220, e202005193. 10.1083/jcb.202005193.

37. Linden, A., Deckers, M., Parfentev, I., Pflanz, R., Homberg, B., Neumann, P., Ficner, R., Rehling, P., and Urlaub, H. (2020). A Cross-linking Mass Spectrometry Approach Defines Protein Interactions in Yeast Mitochondria. Mol Cell Proteomics 19, 1161–1178. 10.1074/mcp.ra120.002028.

38. Schweppe, D.K., Chavez, J.D., Lee, C.F., Caudal, A., Kruse, S.E., Stuppard, R., Marcinek, D.J., Shadel, G.S., Tian, R., and Bruce, J.E. (2017). Mitochondrial protein interactome elucidated by chemical cross-linking mass spectrometry. Proceedings of the National Academy of Sciences 114, 1732–1737. 10.1073/pnas.1617220114.

39. Lee, K., and O’Reilly, F.J. (2023). Cross-linking mass spectrometry for mapping protein complex topologies in situ. Essays Biochem 67, 215–228. 10.1042/ebc20220168.

40. Kolbowski, L., Fischer, L., and Rappsilber, J. (2023). Cleavable crosslinkers redefined by novel MS3-trigger algorithm. Biorxiv, 2023.01.26.525676. 10.1101/2023.01.26.525676.

41. Vogel, F., Bornhovd, C., Neupert, W., and Reichert, A.S. (2006). Dynamic subcompartmentalization of the mitochondrial inner membrane. J. Cell Biol. 175, 237–247. 10.1083/jcb.200605138.

42. Röhricht, H., Przybyla-Toscano, J., Forner, J., Boussardon, C., Keech, O., Rouhier, N., and Meyer, E.H. (2023). Mitochondrial ferredoxin-like is essential for forming complex I-containing supercomplexes in Arabidopsis. Plant Physiol. 10.1093/plphys/kiad040.

43. Brzezinski, P., Moe, A., and Ädelroth, P. (2021). Structure and Mechanism of Respiratory III–IV Supercomplexes in Bioenergetic Membranes. Chem Rev 121, 9644–9673. 10.1021/acs.chemrev.1c00140.

44. Cogliati, S., Calvo, E., Loureiro, M., Guaras, A.M., Nieto-Arellano, R., Garcia-Poyatos, C., Ezkurdia, I., Mercader, N., Vazquez, J., and Enríquez, J.A. (2016). Mechanism of super-assembly of respiratory complexes III and IV. Nature 539, 1–14. 10.1038/nature20157.

45. Pérez-Pérez, R., Lobo-Jarne, T., Milenkovic, D., Mourier, A., Bratic, A., García-Bartolomé, A., Fernández-Vizarra, E., Cadenas, S., Delmiro, A., García-Consuegra, I., et al. (2016). COX7A2L Is a Mitochondrial Complex III Binding Protein that Stabilizes the III2+IV Supercomplex without Affecting Respirasome Formation. Cell Reports 16, 2387–2398. 10.1016/j.celrep.2016.07.081.

46. Lobo-Jarne, T., Nývltová, E., Pérez-Pérez, R., Timón-Gómez, A., Molinié, T., Choi, A., Mourier, A., Fontanesi, F., Ugalde, C., and Barrientos, A. (2018). Human COX7A2L Regulates Complex III Biogenesis and Promotes Supercomplex Organization Remodeling without Affecting Mitochondrial Bioenergetics. Cell Reports 25, 1786–1799.e4. 10.1016/j.celrep.2018.10.058.

47. Ikeda, K., Shiba, S., Horie-Inoue, K., Shimokata, K., and Inoue, S. (2013). A stabilizing factor for mitochondrial respiratory supercomplex assembly regulates energy metabolism in muscle. Nat Commun 4, 2147. 10.1038/ncomms3147.

48. Mourier, A., Matic, S., Ruzzenente, B., Larsson, N.-G., and Milenkovic, D. (2014). The Respiratory Chain Supercomplex Organization Is Independent of COX7a2l Isoforms. Cell metabolism 20, 1069–1075. 10.1016/j.cmet.2014.11.005.

49. Benegiamo, G., Sleiman, M.B., Wohlwend, M., Rodríguez-López, S., Goeminne, L.J.E., Laurila, P.-P., Klevjer, M., Salonen, M.K., Lahti, J., Jha, P., et al. (2022). COX7A2L genetic variants determine cardiorespiratory fitness in mice and human. Nat Metabolism 4, 1336– 1351. 10.1038/s42255-022-00655-0.

50. Granata, C., Caruana, N.J., Botella, J., Jamnick, N.A., Huynh, K., Kuang, J., Janssen, H.A., Reljic, B., Mellett, N.A., Laskowski, A., et al. (2021). High-intensity training induces non-stoichiometric changes in the mitochondrial proteome of human skeletal muscle without reorganisation of respiratory chain content. Nat Commun 12, 7056. 10.1038/s41467-021- 27153-3.

51. Vercellino, I., and Sazanov, L.A. (2021). Structure and assembly of the mammalian mitochondrial supercomplex CIII2CIV. Nature, 1–4. 10.1038/s41586-021-03927-z.

52. Fernández-Vizarra, E., López-Calcerrada, S., Sierra-Magro, A., Pérez-Pérez, R., Formosa, L.E., Hock, D.H., Illescas, M., Peñas, A., Brischigliaro, M., Ding, S., et al. (2022). Two independent respiratory chains adapt OXPHOS performance to glycolytic switch. Cell Metab. 10.1016/j.cmet.2022.09.005.

53. Rappaport, F. (2015). A method aimed at assessing the functional consequences of the supramolecular organization of the respiratory electron transfer chain by time-resolved studies. Methods Mol. Biol. 1241, 95–109. 10.1007/978-1-4939-1875-1_9.

54. Hirst, J. (2018). Open questions: respiratory chain supercomplexes—why are they there and what do they do? Bmc Biol 16, 111. 10.1186/s12915-018-0577-5.

55. Slingsby, C., Wistow, G.J., and Clark, A.R. (2013). Evolution of crystallins for a role in the vertebrate eye lens. Protein Sci. 22, 367–380. 10.1002/pro.2229.

56. Maranzana, E., Barbero, G., Falasca, A.I., Lenaz, G., and Genova, M.L. (2013). Mitochondrial respiratory supercomplex association limits production of reactive oxygen species from complex I. Antioxid. Redox Signal. 19, 1469–1480. 10.1089/ars.2012.4845.

57. Lopez-Fabuel, I., Douce, J.L., Logan, A., James, A.M., Bonvento, G., Murphy, M.P., Almeida, A., and Bolaños, J.P. (2016). Complex I assembly into supercomplexes determines differential mitochondrial ROS production in neurons and astrocytes. Proc. Natl. Acad. Sci. U.S.A. 113, 13063–13068. 10.1073/pnas.1613701113.

58. Protasoni, M., Pérez-Pérez, R., Lobo-Jarne, T., Harbour, M.E., Ding, S., Peñas, A., Diaz, F., Moraes, C.T., Fearnley, I.M., Zeviani, M., et al. (2020). Respiratory supercomplexes act as a platform for complex III-mediated maturation of human mitochondrial complexes I and IV. Embo J 39, e102817. 10.15252/embj.2019102817.

59. Moreno-Lastres, D., Fontanesi, F., García-Consuegra, I., Martín, M.A., Arenas, J., Barrientos, A., and Ugalde, C. (2012). Mitochondrial complex I plays an essential role in human respirasome assembly. Cell metabolism 15, 324–335. 10.1016/j.cmet.2012.01.015.

60. Guerrero-Castillo, S., Baertling, F., Kownatzki, D., Wessels, H.J., Arnold, S., Brandt, U., and Nijtmans, L. (2017). The Assembly Pathway of Mitochondrial Respiratory Chain Complex I. Cell metabolism 25, 1–12. 10.1016/j.cmet.2016.09.002.

61. Letts, J.A., Fiedorczuk, K., Degliesposti, G., Skehel, M., and Sazanov, L.A. (2019). Structures of Respiratory Supercomplex I+III2 Reveal Functional and Conformational Crosstalk. Mol Cell 75, 1131–1146.e6. 10.1016/j.molcel.2019.07.022.

62. Goddard, T.D., Huang, C.C., Meng, E.C., Pettersen, E.F., Couch, G.S., Morris, J.H., and Ferrin, T.E. (2018). UCSF ChimeraX: Meeting modern challenges in visualization and analysis. Protein Sci 27, 14–25. 10.1002/pro.3235.

63. Agip, A.-N.A., Blaza, J.N., Bridges, H.R., Viscomi, C., Rawson, S., Muench, S.P., and Hirst, J. (2018). CryoEM structures of complex I from mouse heart mitochondria in two biochemically-defined states. Nat Struct Mol Biol 25, 548–556. 10.1038/s41594-018-0073-1.

64. Jones, A.J.Y., and Hirst, J. (2013). A spectrophotometric coupled enzyme assay to measure the activity of succinate dehydrogenase☆. Anal Biochem 442, 19–23. 10.1016/j.ab.2013.07.018.

65. Kussmaul, L., and Hirst, J. (2006). The mechanism of superoxide production by NADH:ubiquinone oxidoreductase (complex I) from bovine heart mitochondria. Proceedings of the National Academy of Sciences 103, 7607–7612. 10.1073/pnas.0510977103.

66. Schägger, H., and Jagow, G. von (1991). Blue native electrophoresis for isolation of membrane protein complexes in enzymatically active form. Anal. Biochem. 199, 223–231.

67. Wittig, I., Karas, M., and Schägger, H. (2007). High Resolution Clear Native Electrophoresis for In-gel Functional Assays and Fluorescence Studies of Membrane Protein Complexes*. Mol Cell Proteomics 6, 1215–1225. 10.1074/mcp.m700076-mcp200.

68. Li, X., and Franz, T. (2014). Up to date sample preparation of proteins for mass spectrometric analysis. Arch Physiol Biochem 120, 188–191. 10.3109/13813455.2014.955032.

69. Rappsilber, J., Ishihama, Y., and Mann, M. (2003). Stop and Go Extraction Tips for Matrix-Assisted Laser Desorption/Ionization, Nanoelectrospray, and LC/MS Sample Pretreatment in Proteomics. Anal Chem 75, 663–670. 10.1021/ac026117i.

70. Ritchie, M.E., Phipson, B., Wu, D., Hu, Y., Law, C.W., Shi, W., and Smyth, G.K. (2015). limma powers differential expression analyses for RNA-sequencing and microarray studies. Nucleic Acids Res 43, e47–e47. 10.1093/nar/gkv007.

71. Franz, T., and Li, X. (2012). The OASIS® HLB μElution plate as a one-step platform for manual high-throughput in-gel digestion of proteins and peptide desalting. Proteomics 12, 2487–2492. 10.1002/pmic.201100354.

72. Cox, J., and Mann, M. (2008). MaxQuant enables high peptide identification rates, individualized p.p.b.-range mass accuracies and proteome-wide protein quantification. Nat Biotechnol 26, 1367–1372. 10.1038/nbt.1511.

73. Cox, J., Neuhauser, N., Michalski, A., Scheltema, R.A., Olsen, J.V., and Mann, M. (2011). Andromeda: A Peptide Search Engine Integrated into the MaxQuant Environment. J Proteome Res 10, 1794–1805. 10.1021/pr101065j.

74. Wickham, H. *ggplot2*: *Elegant Graphics for Data Analysis* (Springer-Verlag New York, 2016).

75. R Core Team. R: A Language and Environment for Statistical Computing. (R Foundation for Statistical Computing, 2017).

76. Wessel, D., and Flügge, U.I. (1984). A method for the quantitative recovery of protein in dilute solution in the presence of detergents and lipids. Anal Biochem 138, 141–143. 10.1016/0003-2697(84)90782-6.

77. Klykov, O., Steigenberger, B., Pektaş, S., Fasci, D., Heck, A.J.R., and Scheltema, R.A. (2018). Efficient and robust proteome-wide approaches for cross-linking mass spectrometry. Nat Protoc 13, 2964–2990. 10.1038/s41596-018-0074-x.

78. Jumper, J., Evans, R., Pritzel, A., Green, T., Figurnov, M., Ronneberger, O., Tunyasuvunakool, K., Bates, R., Žídek, A., Potapenko, A., et al. (2021). Highly accurate protein structure prediction with AlphaFold. Nature 596, 583–589. 10.1038/s41586-021-03819-2.

79. Chen, Y.-Y., Chambers, M.C., Li, M., Ham, A.-J.L., Turner, J.L., Zhang, B., and Tabb, D.L. (2013). IDPQuantify: Combining Precursor Intensity with Spectral Counts for Protein and Peptide Quantification. J Proteome Res 12, 4111–4121. 10.1021/pr400438q.

