## Supplementary figures and images for "Preserved respiratory chain capacity and physiology in mice with profoundly reduced levels of mitochondrial respirasomes"

### Supplementary Figure 1

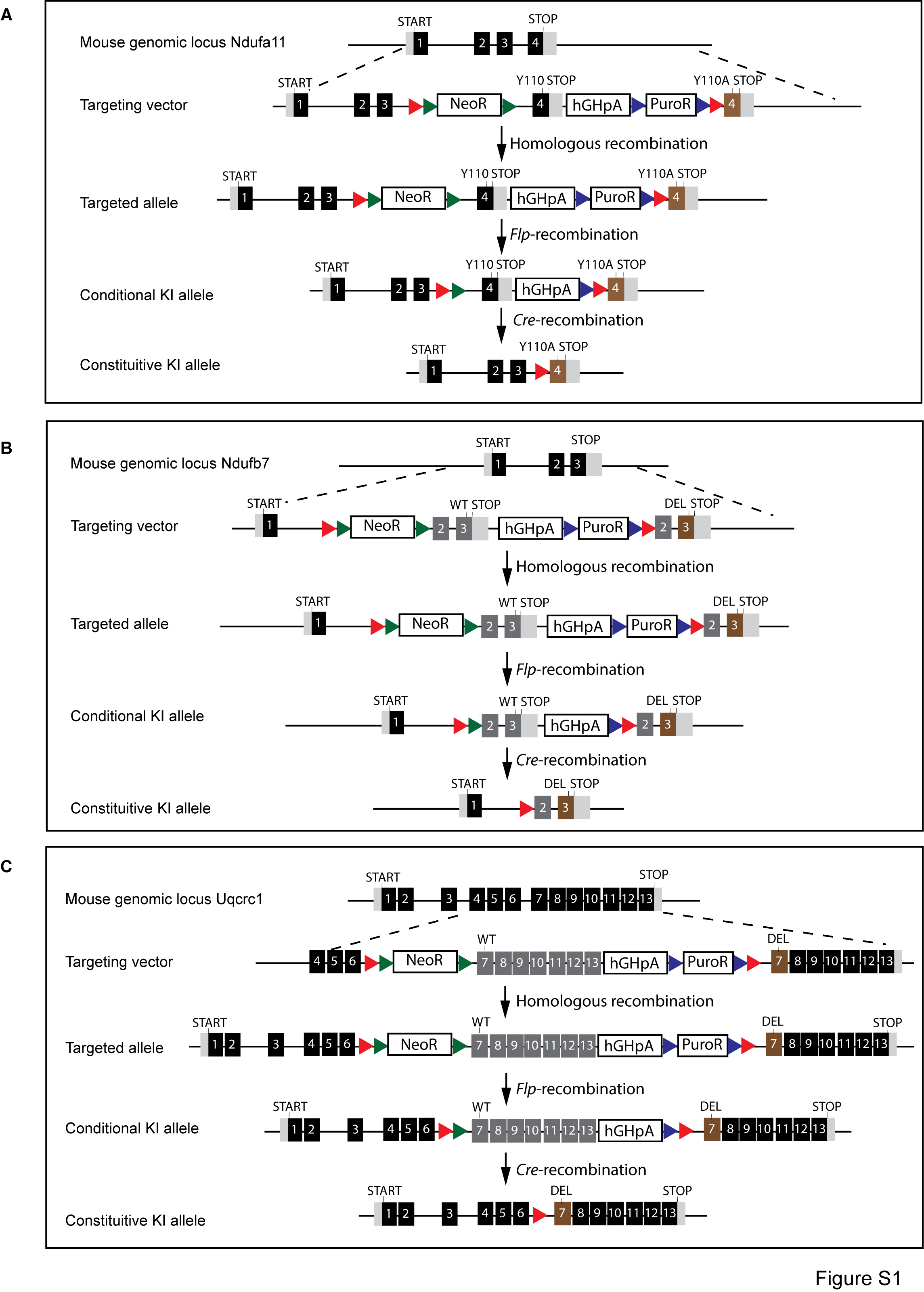

### Supplementary Figure 2

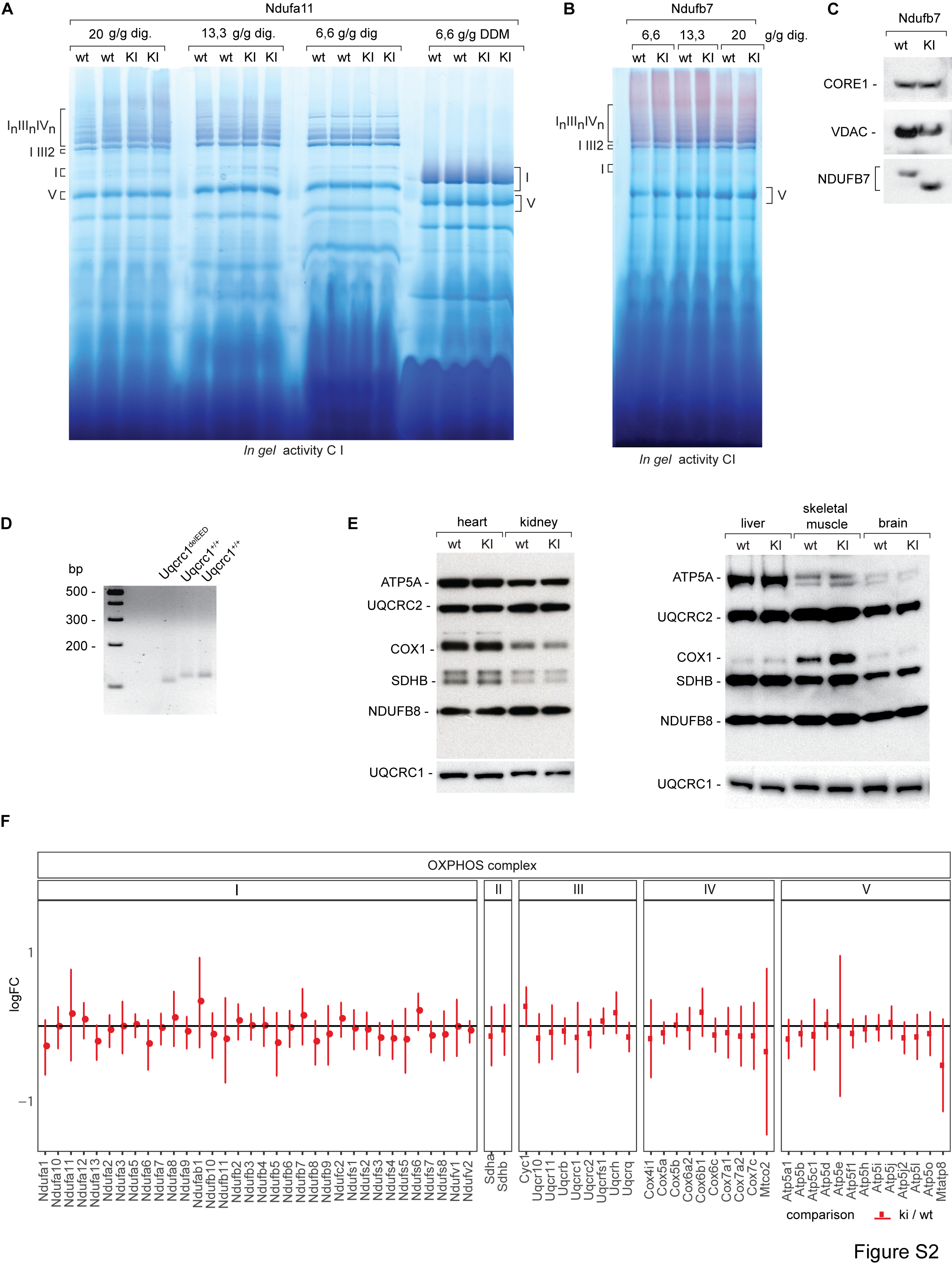

### Supplementary Figure 3

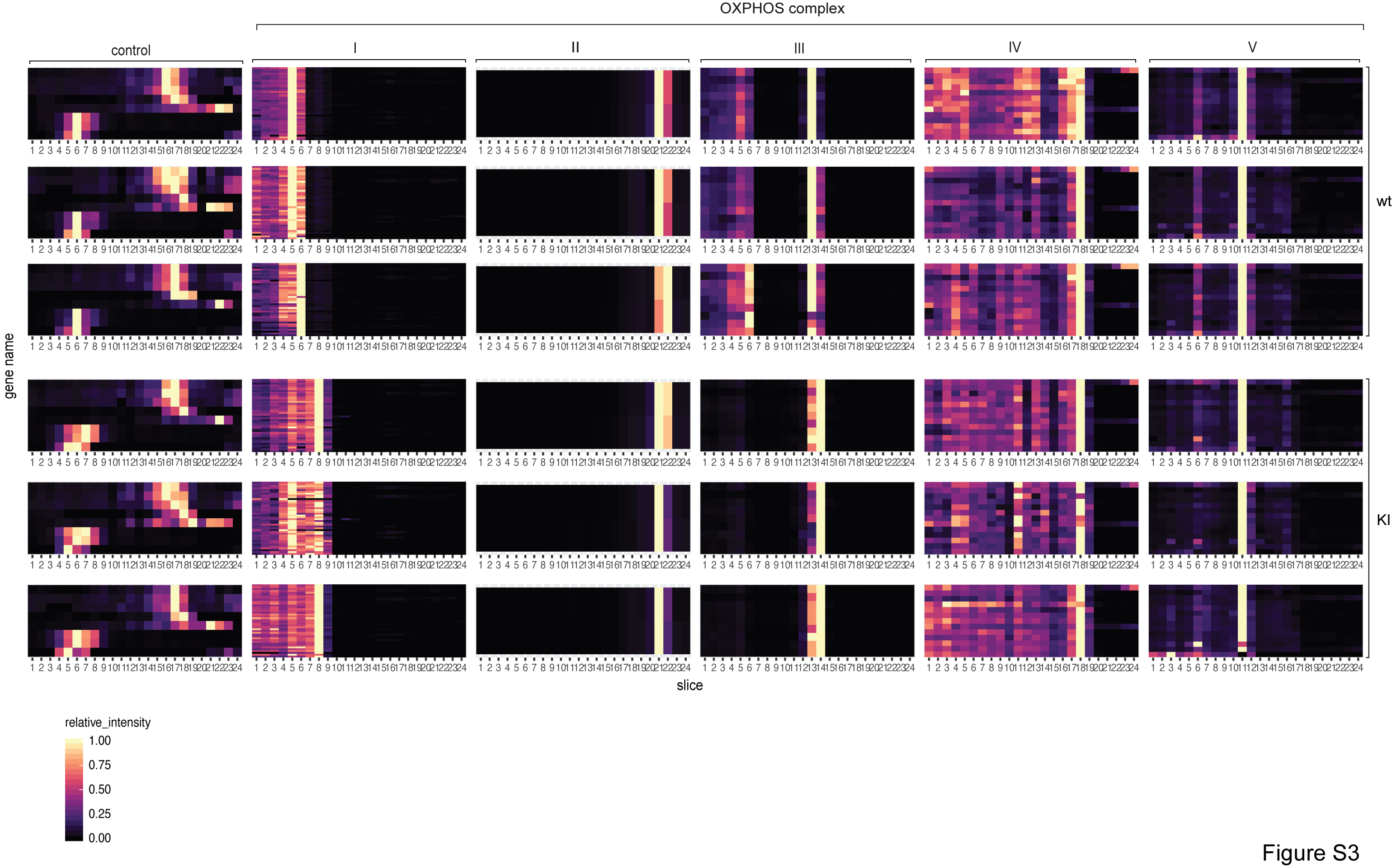

### Supplementary Figure 4

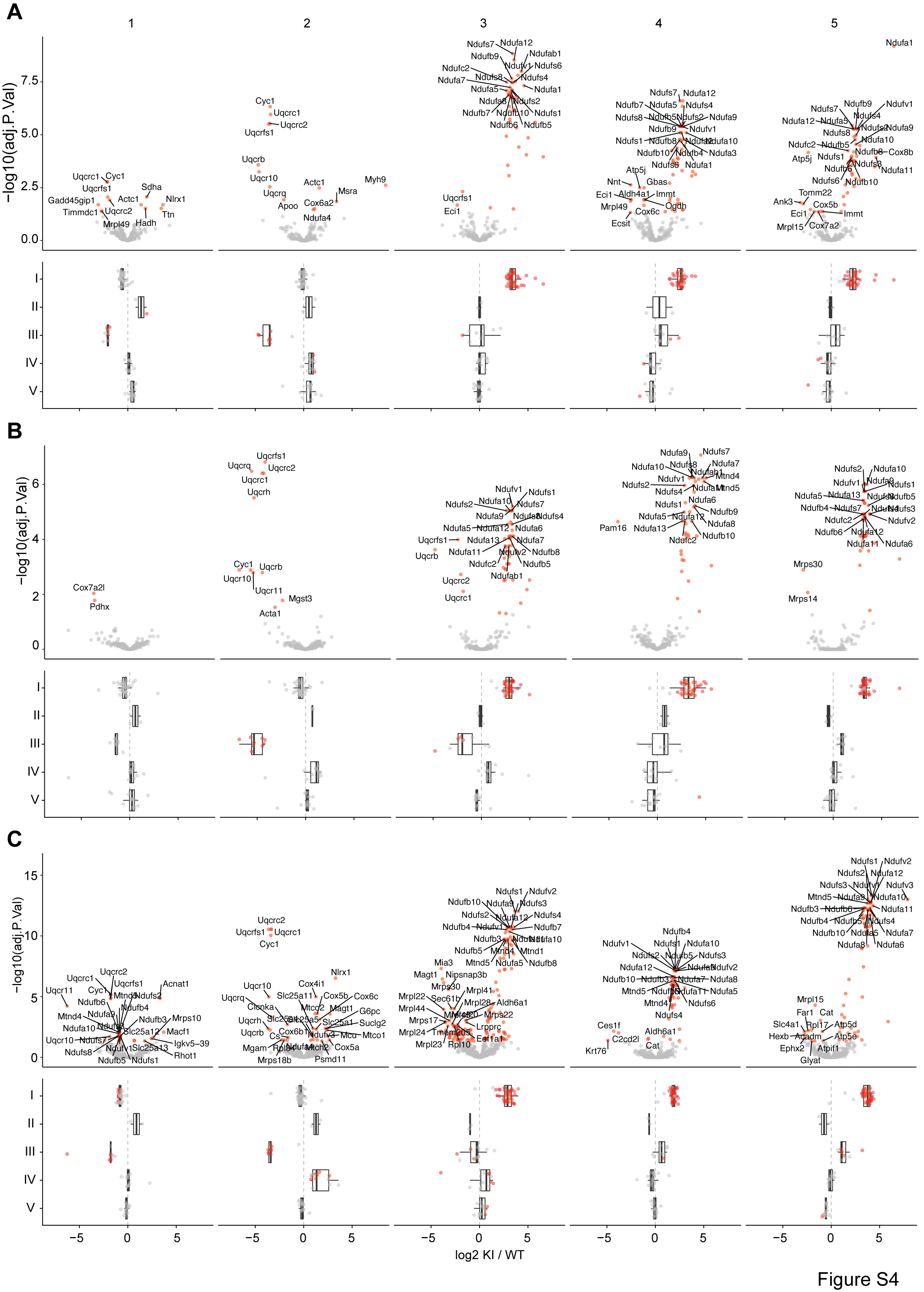

### Supplementary Figure 5

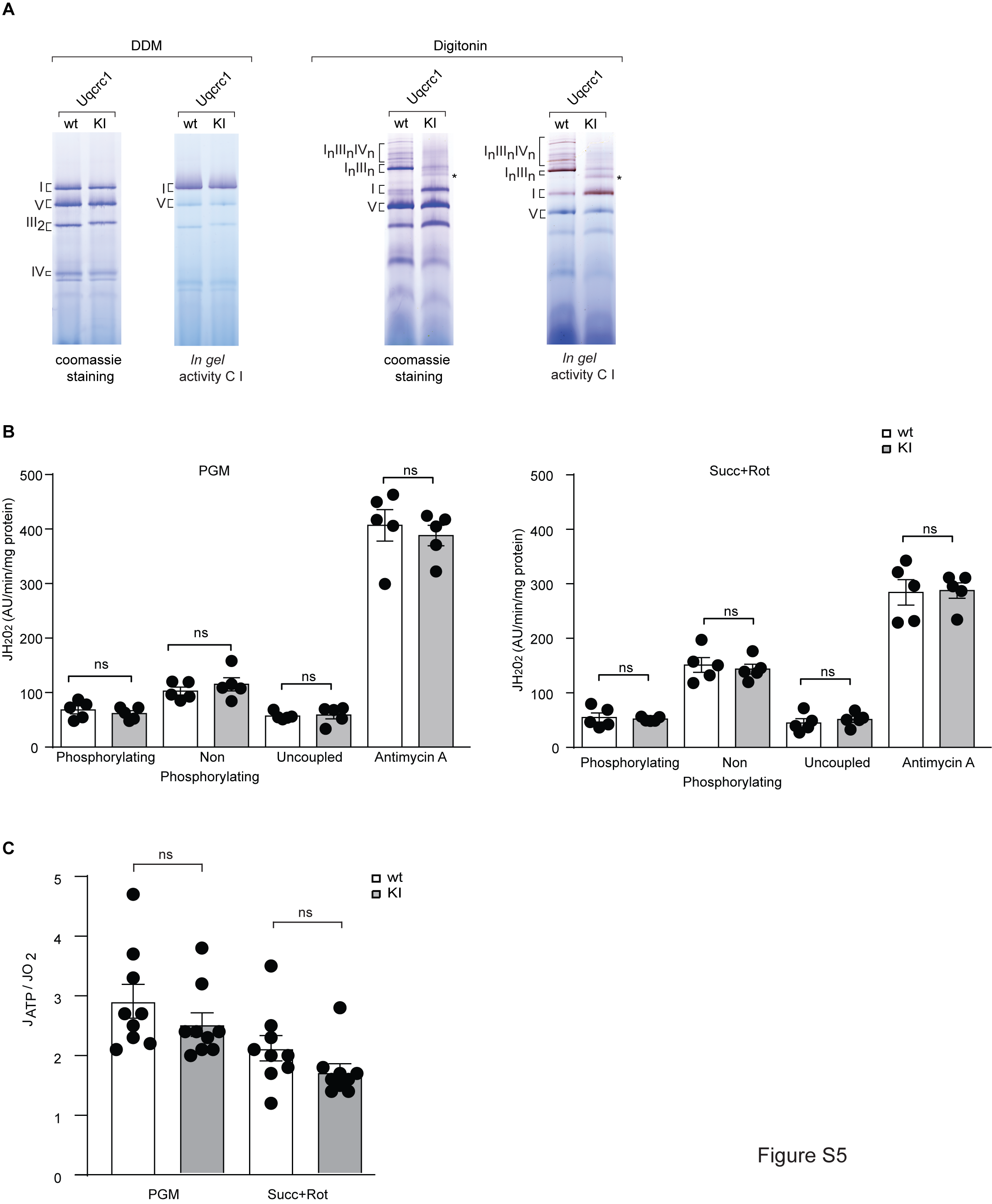

### Supplementary Figure 6

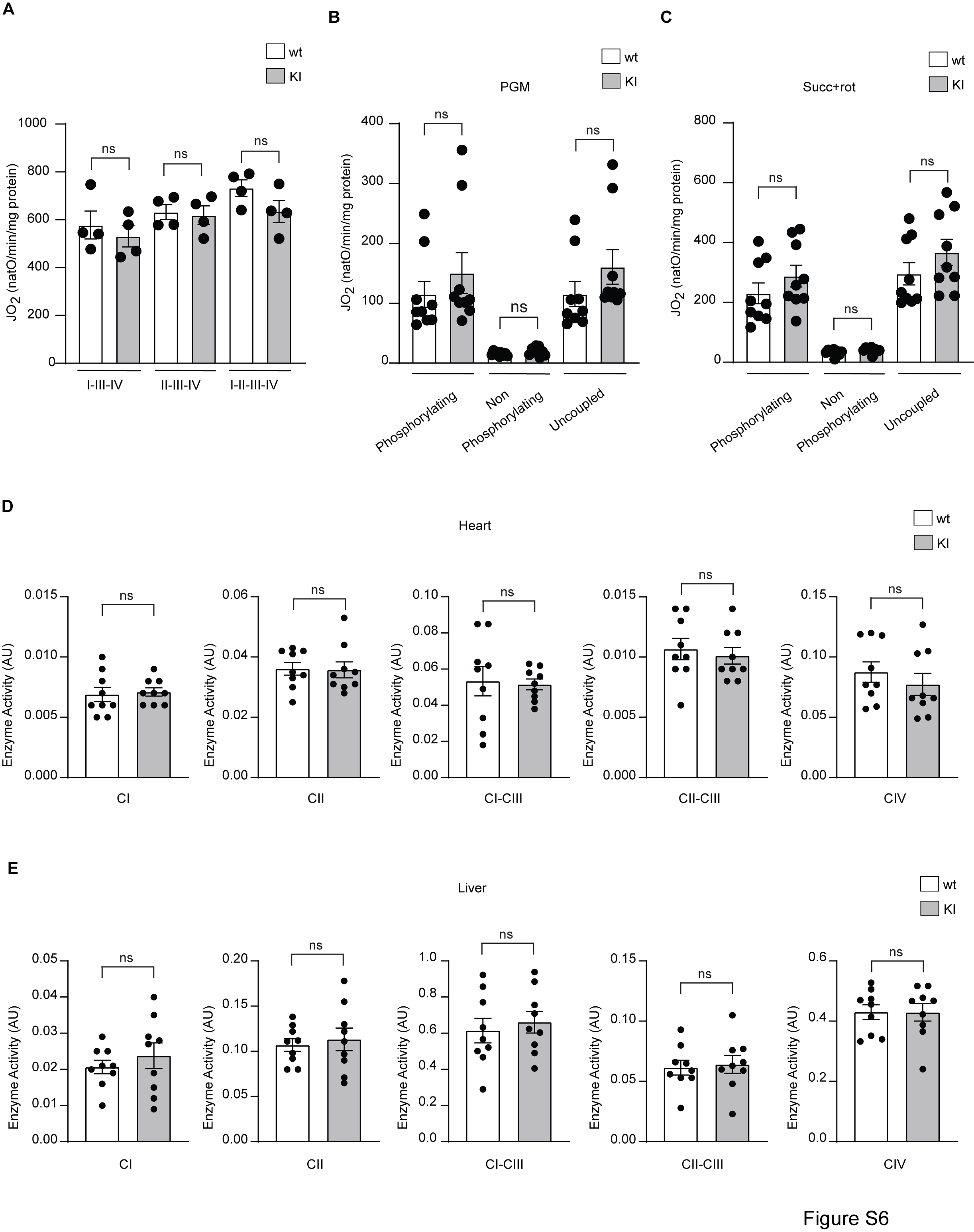

### Supplementary Figure 7

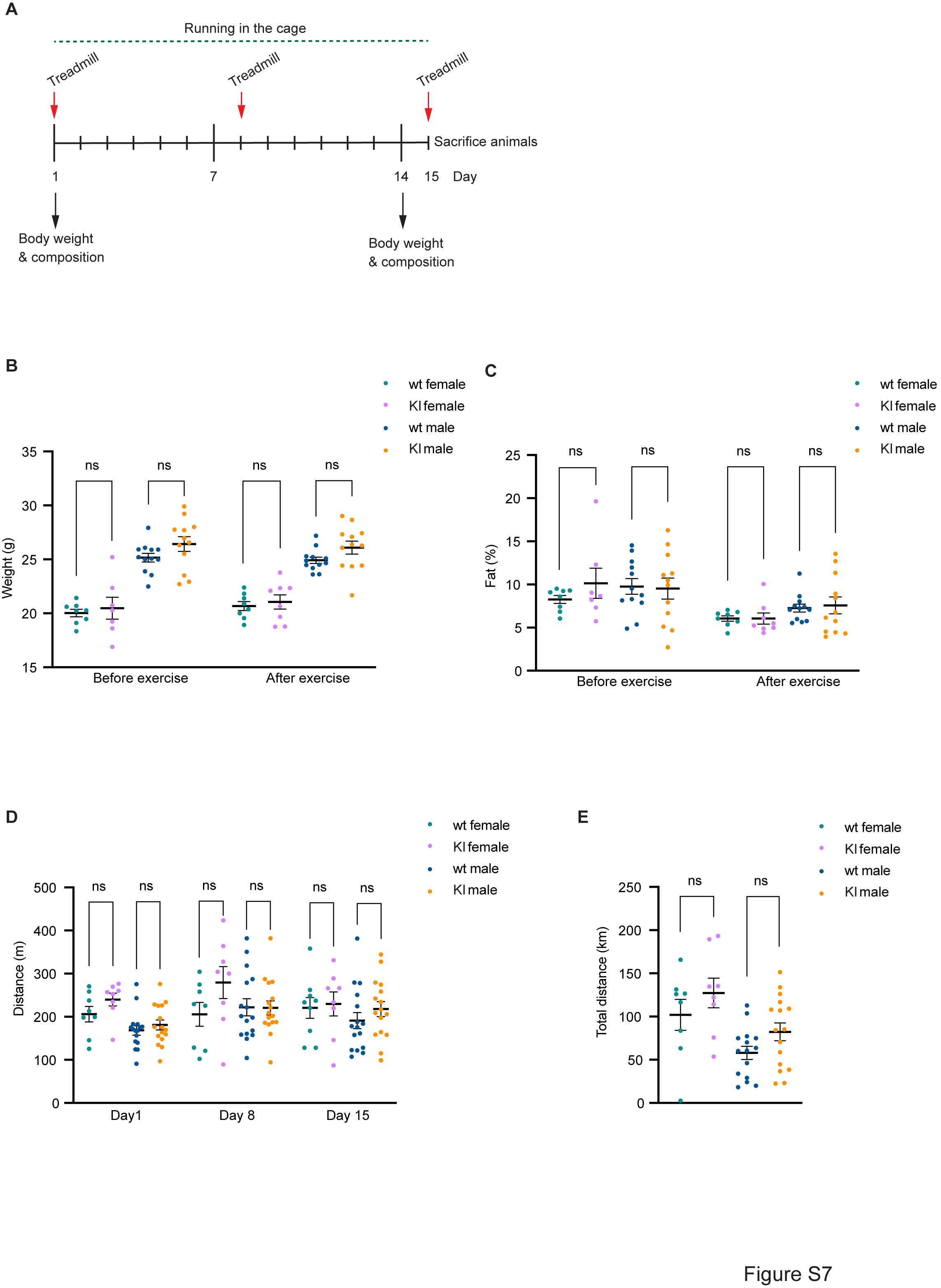

### Supplementary Figure 8

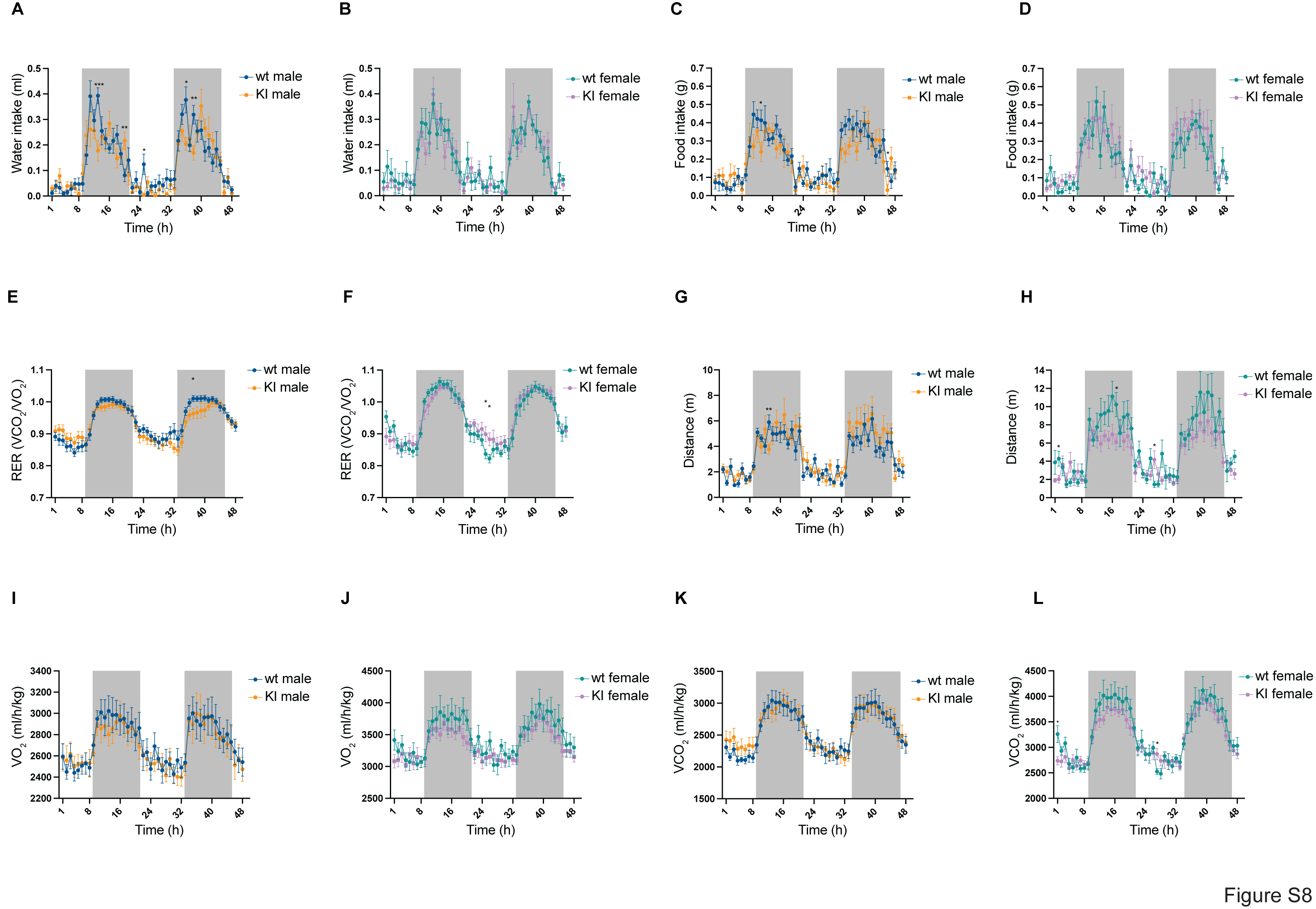

### Supplementary Figure 9

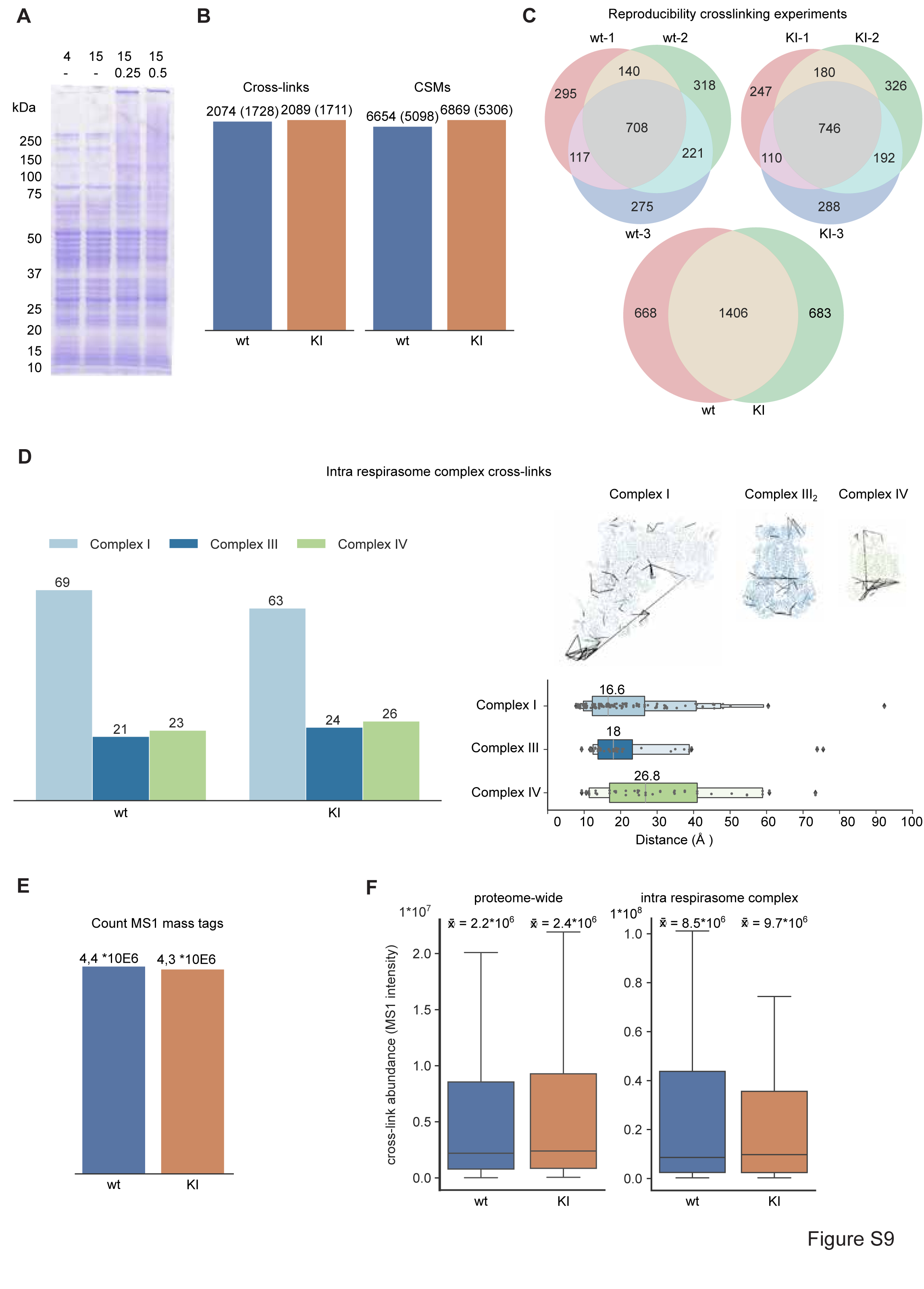
